# Supervised Semantic Similarity

**DOI:** 10.1101/2021.02.16.431402

**Authors:** Rita T. Sousa, Sara Silva, Catia Pesquita

## Abstract

**Background:** Semantic similarity between concepts in knowledge graphs is essential for several bioinformatics applications, including the prediction of protein-protein interactions and the discovery of associations between diseases and genes. Although knowledge graphs describe entities in terms of several perspectives (or semantic aspects), state-of-the-art semantic similarity measures are general-purpose. This can represent a challenge since different use cases for the application of semantic similarity may need different similarity perspectives and ultimately depend on expert knowledge for manual fine-tuning.

**Results:** We present a new approach that uses supervised machine learning to tailor aspect-oriented semantic similarity measures to fit a particular view on biological similarity or relatedness. We implement and evaluate it using different combinations of representative semantic similarity measures and machine learning methods with four biological similarity views: protein-protein interaction, protein function similarity, protein sequence similarity and phenotype-based gene similarity.

**Conclusions:** The results demonstrate that our approach outperforms non-supervised methods, producing semantic similarity models that fit different biological perspectives significantly better than the commonly used manual combinations of semantic aspects.

## Background

The life sciences field has increasingly taken advantage of ontologies to tackle the challenges of managing and analyzing the growing volumes of biomedical data.

Ontologies, in the computer science context, are artifacts that express knowledge about a domain in a shareable and computationally accessible form [1].

They are an explicit specification of a conceptualization in which each element is precisely defined, and the relationships between elements are parameterized or constrained [2]. To enable such a description, ontologies consist of classes and semantic links between the classes as well as restrictions, rules, and axioms. Ontologies often structure their classes, and the relationships between them as a directed acyclic graph, where the classes are nodes and relationships are edges.

Since ontologies are abstractions over reality, they only contain true facts for all entities of a particular type. For that reason, they do not contain entities but instead represent classes only. A semantic annotation is about assigning real-world entities to ontology classes describing them [3]. The ontology data model can be applied to a set of individual entities to create a knowledge graph (KG), where the nodes represent ontology classes and real-world entities, and edges are employed in representing ontology classes’ relations and semantic annotations [4].

In the life sciences, we have witnessed in the last decade not only an increase in the number and size of available ontologies, but also of their relevance in biomedical research [5]. Biomedical ontologies are used for data annotation and management in areas ranging from gene function [6], to biomolecules [7] or phenotypes [8, 9]. There are currently over 800 biomedical ontologies in BioPortal [10].

However, ontologies are also increasingly used to support data analysis and mining[5]. One of the fundamental tasks in this area is measuring the similarity between entities described in an ontology, i.e., semantic similarity[11]. Ontologies allow the description of complex biological phenomena, that are not easily captured in mathematical form. As such, they provide the scaffolding for comparing biological entities at a higher level of complexity by comparing the ontology classes with which they are annotated. There are a wide variety of bioinformatics applications that benefit from using semantic similarity over biomedical ontologies, namely protein-protein interaction (PPI) prediction [12, 13], disease-associated genes identification [14, 15], and drug-drug interaction prediction [16, 17].

The specificity of these data mining tasks is in contrast with the broad domains covered by many biomedical ontologies. Large and successful biomedical ontologies often afford multiple perspectives (or semantic aspects) over the entities it describes. For instance, the Gene Ontology (GO) [6] describes protein function according to three semantic aspects: the *molecular functions* they perform, the *biological processes* they intervene in and the *cellular components* where they are active. In the same way, ChEBI [7] provides information about small chemical entities (e.g., atoms, molecules, ion pairs, radicals, radical ions, complexes, conformers) from three perspectives: the *molecular structure*, the *role* within a biological context or based on the intended use by humans, and the *subatomic particle*. Moreover, it can also be the case that multiple ontologies describe the same real-world entities, each covering a different semantic aspect.

Depending on our viewpoint of the domain or the analytical task for which we want to use semantic similarity, some semantic aspects may be irrelevant for a specific definition of similarity. Consider the following example on comparing proteins according to their function. From a biochemist point of view, two proteins playing the same molecular functions are very similar. However, these proteins can be very different from a physiological perspective if they participate in different biological processes at the whole-organism level. Therefore, depending on our goal, different semantic aspects should be taken into consideration in similarity computation. Selecting which semantic aspects to use and how they should be taken into account usually falls to the domain expert, rendering semantic similarity applications dependent on fine-tuning. This brings us to the challenge of tailoring semantic similarity measures (SSMs) to fit a specific application and biological perspective on similarity. In this work, we present a novel approach that integrates semantic similarity and supervised learning methods to learn semantic similarity models tailored to better capture particular biological similarity views in effect producing a supervised similarity. The proposed approach was implemented using different SSMs coupled with machine learning (ML) methods to elucidate which SSMs are more suitable for different combinations of supervised learning approaches. Since there is no gold standard for the similarity between complex biomedical entities, we take advantage of similarity proxies to train the models and evaluate them. These proxies of similarity rely on objective representations of entities (e.g., gene sequence, domains) and calculate similarity using mathematical expressions or other algorithms (e.g., BLAST-based similarity for sequences).

We evaluate the proposed approach in a set of 21 benchmark datasets [18] that have varying sizes with different semantic annotation characteristics and include data from two biomedical ontologies, GO and Human Phenotype Ontology (HP). These datasets contain four proxies for biomedical entity similarity calculated based on protein sequence similarity, protein function family similarity, protein-protein interactions, and phenotype-based gene similarity. Although these proxy similarities do not provide the broad spectrum comparison that semantic similarity supports, they are known to relate to relevant characteristics of the underlying entities. Our approach is compared with combinations of semantic aspects that emulate expert choices to understand how well the approach captures entity similarity. The results achieved on the benchmark datasets demonstrate the ability of our approach to significantly improve the estimation of similarity between biomedical entities.

## Related Work

A SSM can be defined as a function that estimates the closeness in meaning between two entities. Several SSMs have been proposed with most measures falling in the category of taxonomic semantic similarity (also referred to as ontology-based semantic similarity, or only semantic similarity) [19]. However, graph embeddings, a more recent research direction, can also be used to compute semantic similarity [20, 21, 22].

### Taxonomic Semantic Similarity

Taxonomic semantic similarity compares entities based on the taxonomic relations within the ontology graph [11]. Taxonomic SSMs are generally designed by an expert based on assumptions about how an ontology is used and what should constitute a similarity. They make extensive use of the taxonomical aspect of an ontology, comparing classes based on subclass/superclass relations.

SSMs can be distinguished based on the entities they intend to compare since we can measure the similarity between either ontology classes or real-world entities (annotated with a set of classes). In the case of GO, semantic similarity can be calculated for two ontology classes, for instance, calculating the similarity between two GO classes (e.g., the GO term *protein metabolic process* and the GO term *protein stabilization*); or between two entities each annotated with a set of classes, for instance calculating the similarity between two proteins. Each protein can be annotated with several GO classes so, to assess the similarity between proteins, it is necessary to compare sets of classes rather than single classes.

For class-based semantic similarity, edge-based measures rely on algorithms designed for graph analysis [23, 24]. However, the majority of methods explore the properties of each class involved, typically relying on the information content (IC) of a class, a measure of how informative (or in other words, specific) a class is, and then using it to measure the shared meaning between two classes. IC can be calculated using external data, for instance the frequency of annotations of entities in a corpus [25], or based on intrinsic properties, such as the ontology’s structure [26].

In entity-based semantic similarity, each instance is described with a set of classes which are then processed using one of two approaches: pairwise or groupwise. In pairwise approaches, the semantic similarity is calculated between classes in one set and classes in the other (using class-based measures). In groupwise approaches, the measures can directly compare the sets of classes according to information defined in the ontology, circumventing the need for pairwise comparisons [27, 28]. Purely set-based and vector-based approaches are not common. In vector-based approaches, the sets are compared through their vector representations, with each term corresponding to a dimension, using vector similarity measures.

### Embedding Semantic Similarity

An embedding is a vector representation that maps each node to a lower-dimensional space. The structure of its local graph neighborhood and its graph position are preserved as much as possible. Several methods for building graph embeddings have been proposed [29]. While some focus on exploring the graph facts solely (like translational distance models [30, 31] or semantic matching [32, 33]), others also include additional information, such as entity types, relation paths, axioms and rules, or textual information. More recently, path-based approaches, such as RDF2Vec [34] and Onto2Vec [20], have been proposed by transforming the ontology graph into node sequences. OPA2Vec [21] extends Onto2Vec to, unlike taxonomic SSMs, combine both the formal content of ontologies and the lexical properties of the ontologies. A graph is represented as a set of random walk paths sampled from it, and then natural language methods are applied to the sampled paths for graph embedding.

After employing graph embedding methods, each entity is represented by a vector. It is then possible to compute the graph embedding similarity between two entities by computing the distance of their corresponding vectors in the Euclidean space. If *v*_*i*_ and *v*_*j*_ denote the vector representations of nodes *n*_*i*_ and *n*_*j*_, respectively, the graph embedding similarity *sim*(*n*_*i*_, *n*_*j*_) between nodes *n*_*i*_ and *n*_*j*_ is given by the distance between their vectors *v*_*i*_ and *v*_*j*_ in the Euclidean space. The distance can be computed by the cosine distance:

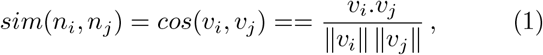

where *v*_*i*_.*v*_*j*_ is the dot product of *v*_*i*_ and *v*_*j*_.

In the GO case, the embedding methods assign to proteins or GO classes a set of points in a low-dimensional space such that similar nodes in the ontology graph correspond to points that are close in the low-dimensional space.

### Machine Learning and Semantic Similarity

More recently, approaches that combine taxonomic semantic similarity with machine learning have been proposed. GARUM [35] is based on a supervised regression algorithm that receives several similarity measures of hierarchy, neighborhood, shared information, and attributes and then predicts a final similarity score. In evoKGsim [36], we have used genetic programming over aspect-oriented semantic similarities to predict protein-protein interactions.

However, the majority of the work that combines ontologies and ML is focused on embeddings. [37] provide an overview of methods that incorporate SSMs and ontology embeddings into ML methods.

## Methods

We have developed a novel approach^[1]^ to learn the similarity between entities represented in KGs (Definition 1) optimized towards a specific similarity proxy. This tailoring is achieved by considering the similarities for different semantic aspects (Definition 2), as opposed to the static SSMs (Definition 3).

### Definition 1

*A* ***KG*** *is created to describe real-world entities using links to ontology classes, represented in a graph. The nodes of the KGs represent ontology classes and entities, and edges are employed in representing ontology classes’ relations and semantic annotations for entities*.

### Definition 2

*A* ***semantic aspect*** *represents a perspective of the representation of KG entities. It can correspond to portions of the graph (e*.*g*., *describing a protein only through the biological process subgraph of the GO) or a given set of property types (e*.*g*., *describing a person only through properties having geographical locations as a range)*.

### Definition 3

*A* ***static SSM*** *calculates values of similarity by processing the KG without additional external input or tailoring to a specific similarity proxy*.

An overview of the approach is shown in Fig 1. The first step consists of identifying the semantic aspects that describe the KG entities. Our approach takes as pre-defined semantic aspects the subgraphs when the KGs have multiple roots (such as GO) or the subgraphs rooted in the classes at a distance of one from the KG root class. As an alternative, semantic aspects can be manually defined. The next step is representing each instance (i.e., a pair of KG entities) according to static KG-based similarities computed for each semantic aspect. The third step in our approach is to select the similarity proxy for which we want to tailor the similarity. The last step is to employ a ML method to learn a supervised semantic similarity. The ML algorithms are used for regression where the expected outputs are the proxy similarity values.

**Figure 1.**
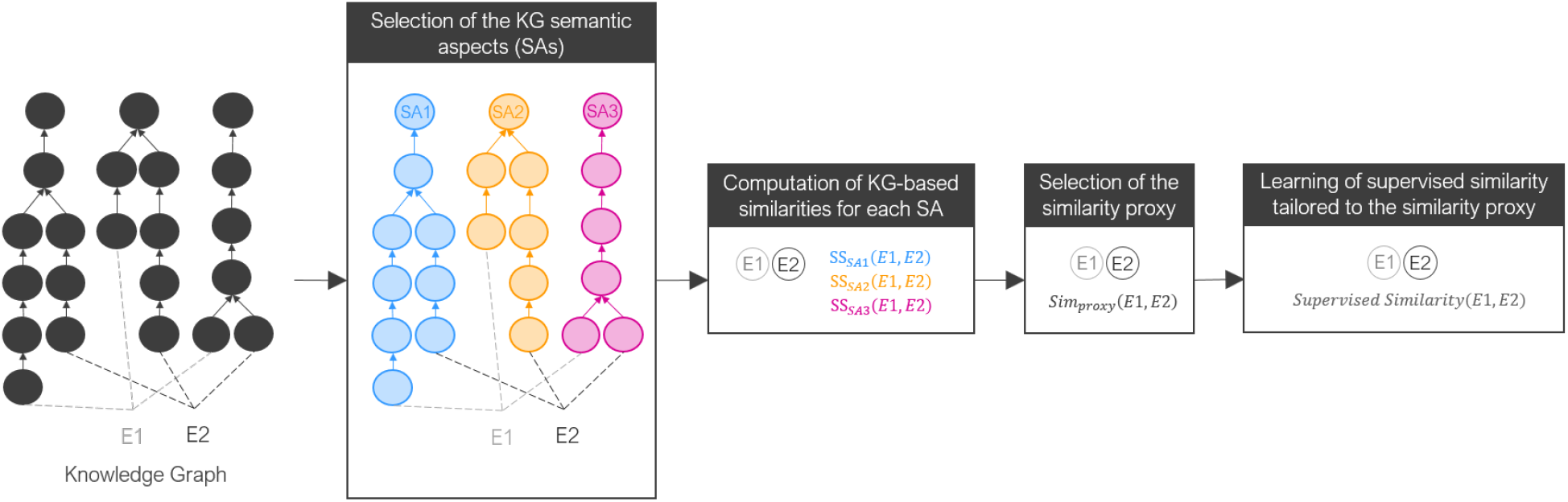
Overview of the proposed approach.

This approach is independent of the semantic aspects, the specific implementation of KG-based similarity and the ML algorithm employed in regression. The following sections present the specific details of the implementation that currently supports five different SSMs (three based on embedding similarity and two based on taxonomic similarity) and eight targeted supervised learning approaches (classical ML approaches and a neural network-based approach).

The approach is evaluated in protein and gene benchmark datasets. For the protein datasets, we consider the GO aspects as semantic aspects. After semantic similarity computations, each instance of the dataset, that represents a protein pair, is characterized by three values corresponding to the semantic similarity between them for the three GO aspects, and a proxy similarity value. The models returned in the second step are then the combinations of the similarity scores of the three GO aspects. For the gene dataset, in addition to the three GO aspects, the similarity is also calculated for the HP phenotypic abnormality subgraph. Therefore, instead of three semantic aspects, we consider four semantic aspects.

### Data

Our approach takes as input an ontology file, an instance annotation file and a list of instance pairs with proxy similarity values. We evaluate our approach using benchmark datasets and two different KGs.

#### Benchmark datasets

The 21 benchmark datasets are presented in [18] and are available online^[2]^ (dated June 2020). These datasets explore four proxy similarities based on protein and gene properties. This resulted in one gene dataset and 16 protein datasets, divided by species, level of annotation completion and similarity proxy, and four additional datasets, combining all species’ protein pairs in the same proxy group. Datasets range in size from 264 individual proteins and 428 pairs to 27 thousand proteins and 158 thousand pairs.

The protein datasets, described in Table 1, are constituted by proteins. Each protein is identified by its UniProt Accession Numbers and annotated with GO classes. Regarding the gene dataset, it has 2026 distinct human genes identified by their Entrez Gene Code and 12000 gene pairs. Each gene is annotated with GO classes and HP classes.

**Table 1.** Number of proteins and pairs for all protein datasets.

|  | PFAM datasets |  | PPI datasets |  |
| --- | --- | --- | --- | --- |
|  | Prots | Pairs | Prots | Pairs |
| DM1 | 7494 | 53797 | 481 | 397 |
| DM3 | 5810 | 52460 | 335 | 270 |
| EC1 | 1250 | 4623 | 371 | 738 |
| EC3 | 748 | 1813 | 264 | 428 |
| SC1 | 4783 | 42192 | 3874 | 34772 |
| SC3 | 3660 | 30747 | 2959 | 21577 |
| HS1 | 13604 | 60176 | 7644 | 44677 |
| HS3 | 12487 | 60163 | 7149 | 42204 |
| ALL1 | 27131 | 158512 | 12370 | 80584 |
| ALL3 | 22705 | 142736 | 10707 | 64479 |

Table 2 shows the similarity approaches employed. In the PFAM datasets, two proxies of protein similarity based on their biological properties were employed: sequence similarity and PFAM similarity. In PPI protein datasets, two similarity proxies were also employed: sequence similarity and protein-protein interactions. Concerning the gene benchmark dataset, the proxy similarity is based on phenotypic series.

**Table 2.** Available similarity proxies for each dataset type.

| Dataset Type | Similarity Proxies |
| --- | --- |
| PFAM | $\text{Sim}_{\text{seq}}$ and $\text{Sim}_{\text{PFAM}}$ |
| PPI | $\text{Sim}_{\text{seq}}$ and PPI |
| Gene | $\text{Sim}_{\text{PS}}$ |

- ***Sequence similarity*** (Sim_seq_) measures the relationship between two sequences and it establishes the likelihood for sequence homology. We infer homology (i.e., common evolutionary ancestry) when two sequences share more similarity than would be expected by chance. A sequence similarity value is aimed to approximate the evolutionary distance between proteins.
- ***PFAM similarity*** (Sim_PFAM_) is computed by comparing the functional regions (commonly termed domains) that exist in each protein sequence. Protein functional domains were extracted from the PFAM [38]. Since protein domains typically correspond to functional sites of a protein, determining similarity between domains can help to define protein function.
- ***Protein-protein interaction*** (PPI) has a binary representation: 1 if the proteins interact, 0 otherwise. Two proteins are considered to be similar if they interact. PPIs are responsible for many critical functions in biology and are highly relevant to disease states.
- ***Phenotypic series similarity*** (Sim_PS_) is based on OMIM’s Phenotypic Series [39], which are groups of identical or similar phenotypes and their associated genes. Phenotypic similarity reflects the similarity between genes and can help to find biological modules of functionally related genes.

#### Gene ontology knowledge graph

GO [6] is the most widely used biological ontology. It defines the universe of classes, also called “GO terms”, associated with gene product (proteins or RNA) functions and how these functions are related with each other with respect to these three aspects: (i) molecular function (MF), the activities that occur at the molecular level performed by the gene product; (ii) biological process (BP), the larger process in which the gene product is active;; (iii) cellular component (CC), the cellular compartments in which the gene product performs a function. Fig 2 shows a small fraction of the GO and annotated proteins.

**Figure 2.**
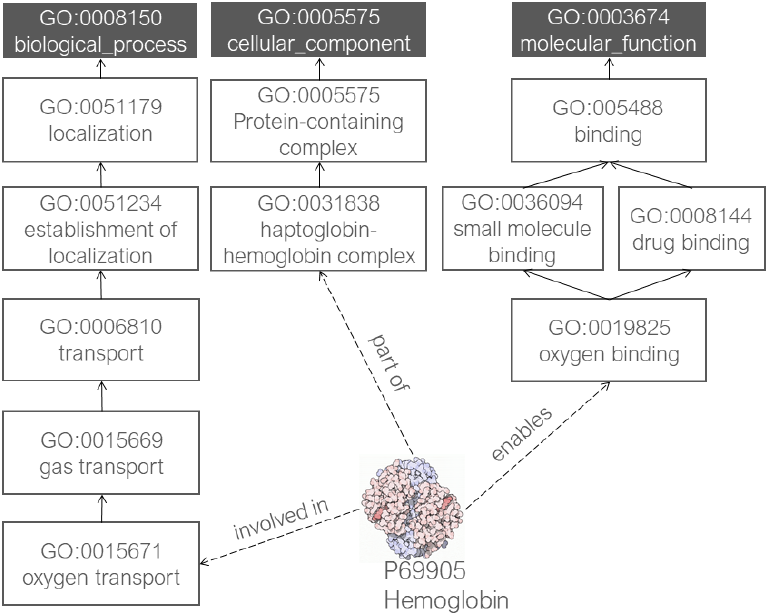
Graph representation of part of GO and GO Annotations.

We built the GO KG with GO, proteins as instances, and GO annotations. Therefore, the nodes of the GO KG represent proteins or GO classes. The KG edges represent relationships between the GO classes or links between proteins annotated with GO classes. The most commonly used relationships between GO classes are *is*_*a*_; *part*_*o*_*f* ; *has*_*p*_*art*; *regulates*; *negatively*_*r*_*egulates* and *positively*_*r*_*egulates*. In this work, the GO KG, with its three semantic aspects (BP, CC and MF), is used to compute the similarity between two proteins for the protein datasets and between two genes for the gene dataset.

#### Human phenotype knowledge graph

The HP [8] contains about terms describing phenotypic abnormalities found in human hereditary diseases. The HP is organized as independent subontologies that cover different categories: “Phenotypic abnormality”, “Mode of inheritance”, “Clinical course”, “Clinical modifier” and “Frequency”. Since the sub-ontology “Phenotypic Abnormality” is the ontology branch that describes the phenotypes the gene is associated with, the HP KG is composed by this subontology and HP annotations. An HP annotation associates a specific gene with a specific HP class. Fig 3 shows the HP and HP annotations.

**Figure 3.**
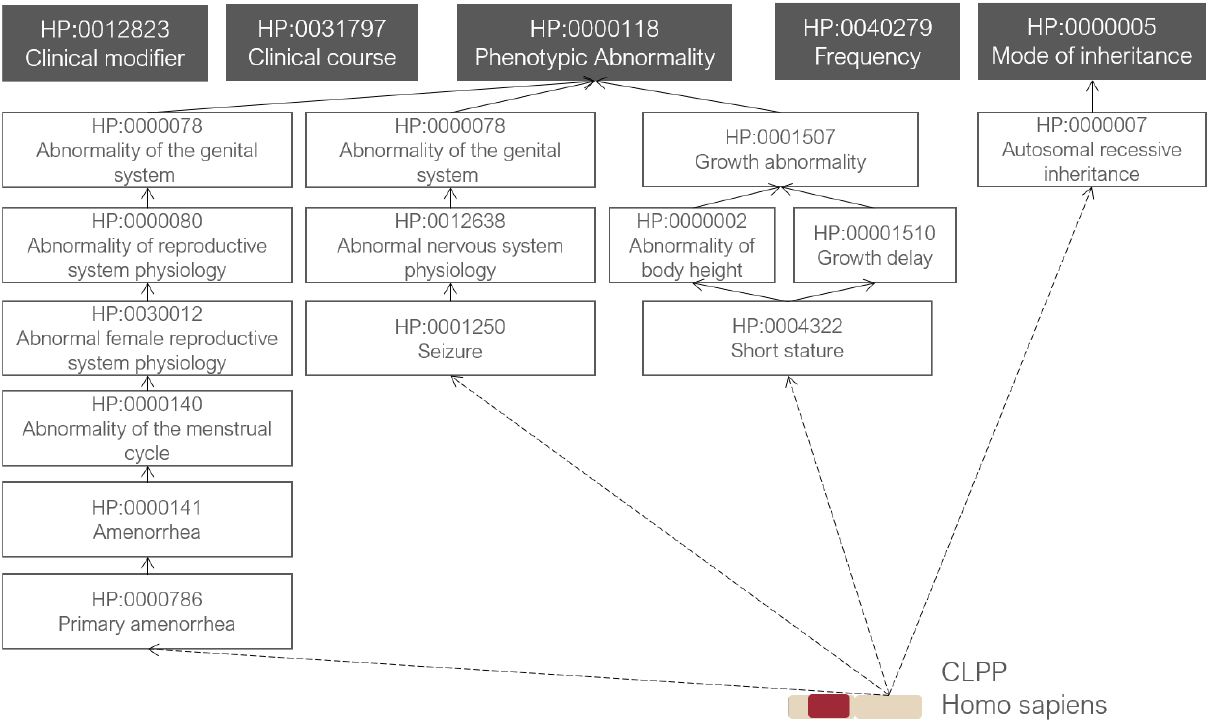
Graph representation of part of HP and HP Annotations.

We consider the HP, genes and associated HP annotations to compose the HP KG. The nodes of HP KG are HP classes or genes. The edges represent ontology relations or links between genes and HP classes via their annotations. In this work, the HP KG is used to compute the semantic similarity between two genes based on the phenotypes that describe them.

### Static similarity computation

The following subsections present the specific details of the five different KG-based SSMs: two based on taxonomic similarity and three based on embeddings.

#### Taxonomic semantic similarity

The taxonomic semantic similarity is calculated using two state-of-the-art measures, derived by combining one IC approach (IC_Seco_) with one of two set similarity measures (ResnikBMA, SimGIC). These were selected by their high performance in the biomedical domain [40].

IC_Seco_ is a structure-based approach proposed by [26] based on the number of direct and indirect descendants and given by

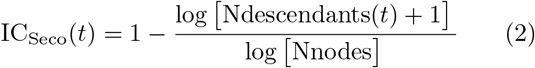

where Ndescendants(*t*) is the number of indirect and direct descendants from term *t* (including term *t*) and Nnodes is the total number of concepts in the ontology.

***ResnikBMA*** is based on the class-based measure proposed by Resnik [25] in which the similarity between two classes corresponds to the IC of their most informative common ancestor and given by:

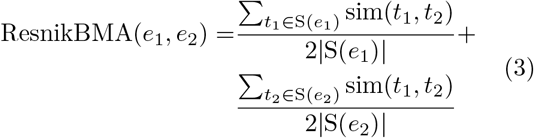

where S(*e*_*i*_) is the set of annotations for entity *e*_*i*_ and sim(*t*_1_, *t*_2_) is the semantic similarity between class *t*_1_ and class *t*_2_ and is defined as:

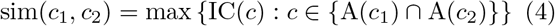

where A(*c*_*i*_) is the set of ancestors of *c*_*i*_.

***SimGIC*** is a groupwise approach proposed by [27], based on a Jaccard index in which each term is weighted by its IC and given by

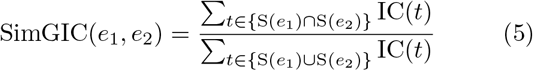

where S(*e*_*i*_) is the set of annotations (direct and inherited) for entity *e*_*i*_.

The Semantic Measures Library 0.9.1 [41] was used to compute the taxonomic semantic similarity.

#### Graph embedding similarity

We employ three graph embedding approaches, namely RDF2Vec, TransE, and distMult, using an RDF2Vec python implementation^[3]^ and the OpenKE library^[4]^.

These approaches were selected because they are representative of the main types of graph embedding techniques.

- ***RDF2Vec*** [34] is a path-based approach adapted to RDF graphs. In RDF2Vec, edge direction is taken into account, enriching the semantics of the learning approach, and neural language models are employed over random walks on the RDF graph to produce the embeddings.
- ***TransE*** [30] is the most representative translational distance embedding approach that exploits distance-based scoring functions. In translational distance models, each fact represents the distance between the two entities, usually after a translation carried out by the relations.
- ***distMult*** [32] is a semantic matching approach that exploits similarity-based scoring functions by matching latent semantics of entities and relations embodied in their vector space representations. DistMult takes the inherent structure of relations into account by employing the tensor factorization.

We generate protein or gene graph embeddings for each semantic aspect using these approaches^[5]^ and then, to compute the graph embeddings similarities, we employ cosine similarity between the vectors representing each entity in the pair.

### Supervised similarity computation

Our approach combines the semantic similarities computed for each semantic aspect and returns a supervised similarity. The aggregation function is computed by a supervised regression algorithm. Therefore, each regressor receives the similarity values for each semantic aspect as input features (independent variables) and a similarity proxy value as the expected output (dependent variable), and returns an aggregated similarity score as the predicted output. We employ eight well-known classes of ML models to train regressors using scikit-learn 21.3 [42] library: linear regression (LR), bayesian ridge (BR), *K*-nearest neighbor (KNN), genetic programming (GP), decision tree (DT), random forest (RF), XGBoost (XGB), and multi-layer perception (MLP).

These algorithms are selected as representative of different types of ML methods. LR [43] assumes there is a linear relationship between the independent and dependent variables. BR [44] is also a linear model but uses the Bayes theorem to find the posterior distribution over all parameters. KNN [45] explores the feature space and reaches a prediction for each sample based on the expected outputs of its neighbors. In DT [46], trees are constructed by beginning with the root node that contains the whole learning sample and then splitting a node into two child nodes repeatedly. The basic idea of tree growth is to choose, among all the possible splits at each node, a split whose resulting child nodes are the “purest”. GP [47] is an evolutionary computation technique inspired by Darwinian natural selection and Mendelian genetics. GP tries to optimize a combination of variable and operators/functions. MLP [48] is a class of feedforward artificial neural networks that learn non-linear functions for regression through back-propagation of errors. At last, RF [49] and XGB [50] are ensemble methods. The goal of ensemble methods is to combine the decisions from multiple models to improve the overall performance. RF builds several estimators independently and then combines their predictions through a voting scheme, while XGB builds base estimators sequentially and tries to reduce the bias of the combined estimator through gradient boosting.

Except for GP, XGB and RF, the parameters are used with the default scikit-learn values. For running GP, we use gplearn 3.0^[6]^, a freely available package that runs with the scikit-learn library with the parameters proposed in [36]. For XGB, we use the XGBoost 1.1.1 package^[7]^, with the values of some parameters altered to maximize the performance of the model, through grid search. For RF, using scikit-learn, we also optimize some parameters^[8]^.

### Supervised similarity evaluation

The focus of our evaluation approach is to assess the ability of our approach to improve semantic similarity computations, avoiding the need for expert knowledge. For each combination of an SSM with an ML algorithm, we compute the Pearson’s correlation coefficient between the obtained supervised similarity (predicted values) and the respective similarity proxies (expected values).

For cross-validation, each dataset is split into ten folds. The same ten folds are used throughout all the experiments. For each fold, we take that fold as the test set and the remaining nine folds as the training set. Each ML algorithm learns on the training set and outputs its predictions for the test set, where the Pearson correlation coefficient is calculated. The results we report are the median and the interquartile range (IQR) of the ten Pearson correlation coefficients calculated on the ten folds.

We compute the static similarity for each semantic aspect and use, as baselines, the single aspect similarities and two well-known strategies for combining the single aspect scores, the average and maximum. By comparing these baselines to the supervised approaches, we aim at investigating the ability of ML methods to learn combinations of semantic aspects that improve the calculation of similarity.

## Results and Discussion

### Static similarity

Before performing the comparative evaluation, we investigate the behavior of the five similarity-based semantic measures employed. Tables 3 and 4 show the Pearson correlation coefficient between the static similarity and the proxy similarity using different SSMs for the gene and protein datasets, respectively. For the sake of simplicity, Table 3 presents only the correlation for the protein datasets combining all species’ protein pairs in the same group proxy, including proteins with at least one annotation in each GO aspect (ALL1). Supplementary File provides the results for the remaining protein datasets.

**Table 3.** Pearson correlation coefficient between static semantic similarity and similarity proxies for the protein datasets with one level of annotation.

| Proxy | Dataset | SSM | Static Similarity |  |  |  |  |
| --- | --- | --- | --- | --- | --- | --- | --- |
|  |  |  | BP | CC | MF | AVG | MAX |
| Sim<sub>seq</sub> | PFAM | ResnikBMA | 0.528 | 0.373 | 0.291 | 0.481 | 0.399 |
|  |  | SimGIC | 0.552 | 0.406 | 0.415 | 0.547 | 0.406 |
|  |  | RDF2Vec | 0.540 | 0.437 | 0.419 | 0.544 | 0.457 |
|  |  | TransE | -0.020 | -0.017 | -0.001 | -0.021 | -0.011 |
|  |  | distMult | 0.398 | 0.236 | 0.322 | 0.467 | 0.429 |
|  | PPI | ResnikBMA | 0.258 | 0.222 | 0.326 | 0.323 | 0.250 |
|  |  | SimGIC | 0.317 | 0.257 | 0.370 | 0.380 | 0.280 |
|  |  | RDF2Vec | 0.274 | 0.237 | 0.297 | 0.316 | 0.268 |
|  |  | TransE | 0.355 | 0.352 | 0.364 | 0.498 | 0.401 |
|  |  | distMult | 0.277 | 0.239 | 0.202 | 0.369 | 0.310 |
| Sim<sub>PFAM</sub> | PFAM | ResnikBMA | 0.448 | 0.370 | 0.456 | 0.525 | 0.500 |
|  |  | SimGIC | 0.494 | 0.451 | 0.591 | 0.621 | 0.604 |
|  |  | RDF2Vec | 0.524 | 0.466 | 0.619 | 0.627 | 0.623 |
|  |  | TransE | -0.032 | -0.031 | -0.011 | -0.039 | -0.027 |
|  |  | distMult | 0.414 | 0.254 | 0.388 | 0.516 | 0.457 |
| PPI | PPI | ResnikBMA | 0.545 | 0.486 | 0.353 | 0.569 | 0.545 |
|  |  | SimGIC | 0.486 | 0.428 | 0.318 | 0.496 | 0.458 |
|  |  | RDF2Vec | 0.457 | 0.404 | 0.353 | 0.477 | 0.440 |
|  |  | TransE | 0.031 | 0.087 | 0.035 | 0.071 | 0.076 |
|  |  | distMult | 0.452 | 0.244 | 0.015 | 0.388 | 0.396 |

**Table 4.** Pearson correlation coefficient between static semantic similarity and similarity proxies for the human gene dataset.

| Proxy | SSM | Static Similarity |  |  |  |  |  |
| --- | --- | --- | --- | --- | --- | --- | --- |
|  |  | HP | BP | CC | MF | AVG | MAX |
| Sim<sub>PS</sub> | ResnikBMA | 0.601 | 0.210 | 0.142 | 0.055 | 0.413 | 0.552 |
|  | SimGIC | 0.489 | 0.205 | 0.158 | 0.095 | 0.399 | 0.429 |
|  | RDF2Vec | 0.526 | 0.230 | 0.182 | 0.123 | 0.396 | 0.351 |
|  | TransE | 0.042 | -0.019 | -0.012 | -0.019 | -0.004 | 0.016 |
|  | distMult | 0.015 | 0.184 | 0.105 | 0.041 | 0.179 | 0.182 |

For most datasets, the behavior of each SSM is consistent. Comparing the two taxonomic semantic similarity approaches, we verify that, in most cases, the maximum correlation is achieved when the ResnikBMA approach is used. Regarding the graph embedding approaches, TransE has performed worse than the other embedding methods. These differences are not unexpected, since the methods that put more emphasis on local neighborhoods, such as translational distance approaches, are less suitable since they fail to capture longer-distance relations. This is relevant when most of the information to be processed is represented in the ontology portion of the KG, where taxonomic relations play an essential role. For that reason, in the following sections, the results obtained with TransE were excluded. distMult, a semantic matching method, is the second-best class of embeddings. Finally, RDF2Vec, a path-based approach, can capture taxonomic (longer-distance) relations which translates into a broader representation of the entities, achieving better results than the other embedding methods in most experiments.

When comparing the two types of semantic similarity, taxonomic similarity performs well across many evaluations and, in the majority of the datasets, has better performance than embedding similarity. The initial assumption was that embedding similarity could potentially outperform taxonomic similarity since semantic similarity is limited to the taxonomic relations within the ontology. In contrast, embeddings take into account all types of relations and, therefore, the embedding representations could, in principle, be more informative. However, the ability of taxonomic similarity to take into account class specificity may give it the advantage over embedding similarity to estimate similarity more accurately. Besides, taxonomic similarity measures are usually hand-crafted, providing human interpretable results for further analysis. On the contrary, embeddings methods describe an entity as a numerical vector and, most of the times, are not interpretable since it is not possible to obtain an explanation for the results.

It is also important to point out the differences between semantic aspects. These differences depend on the similarity proxy we are considering. For the sequence proxy, the differences between semantic aspects are not relevant, and there is no semantic aspect clearly superior to others. Previous works [51] already suggested the absence of a strong correlation between sequence and semantic similarities since there is a large number of proteins with low sequence similarity and high semantic similarity. Concerning the PPI proxy, proteins that interact in a cell are expected to participate in similar cellular locations and processes. As expected, the results indicate that using only the semantic similarity for MF provides worse results compared to the other single aspects. In opposition, for the PFAM proxy, we verify that the MF is a relevant semantic aspect. The more functional (or PFAM) domains two proteins share, the more likely it will be to share molecular functions since these domains are usually responsible by assigning functions to proteins. For the gene dataset, the HP semantic aspect achieves better results compared to the GO semantic aspects. These results were also expected since the more phenotypic series two genes are associated with, the more likely it is that they share HP classes. Regarding static combination approaches, in most cases, they achieve better results than the single aspects, with the average combination outperforming the maximum.

### Supervised similarity

The similarity proxies reflecting different biological features allow us to use ML algorithms to learn a supervised similarity towards a viewpoint of the domain.

We employ eight representative ML methods, including classical, ensemble and neural network based methods: linear regression (LR), bayesian ridge (BR), *K*-nearest neighbor (KNN), genetic programming (GP), decision tree (DT), random forest (RF), XGBoost (XGB), and multi-layer perception (MLP).

Figs 4 to 8 contain the heat maps depicting the median Pearson correlation coefficient between the similarity proxies and supervised similarity obtained with different ML methods and SSMs for each similarity proxy. In some cases, the results obtained with GP are not shown because the GP model only contains constants and, therefore, the Pearson correlation coefficient is undefined (division by zero). To better compare the eight ML algorithms, we also generated radar charts (Figs 9 to 13) showing the median Pearson correlation coefficient between supervised similarity and each similarity proxy. Radar charts reveal which ML algorithms combined with different SSMs are scoring high or low within a dataset. In each radar plot, the ML algorithms are represented by different colors^[9]^, and the SSMs are represented on different axes.

**Figure 4.**
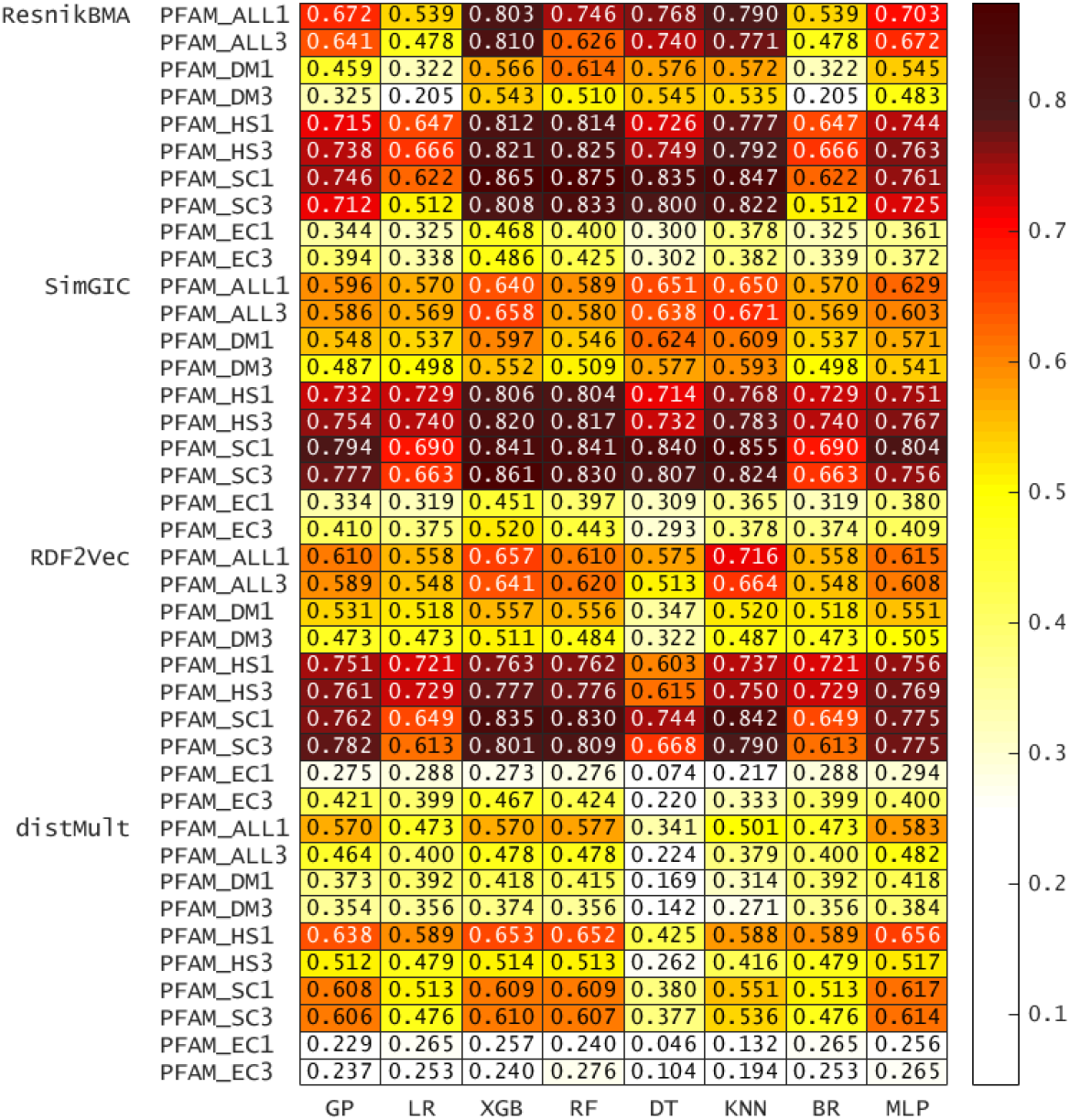
Heat map representing the median Pearson’s correlation coefficient using sequence proxy for each PFAM dataset.

**Figure 5.**
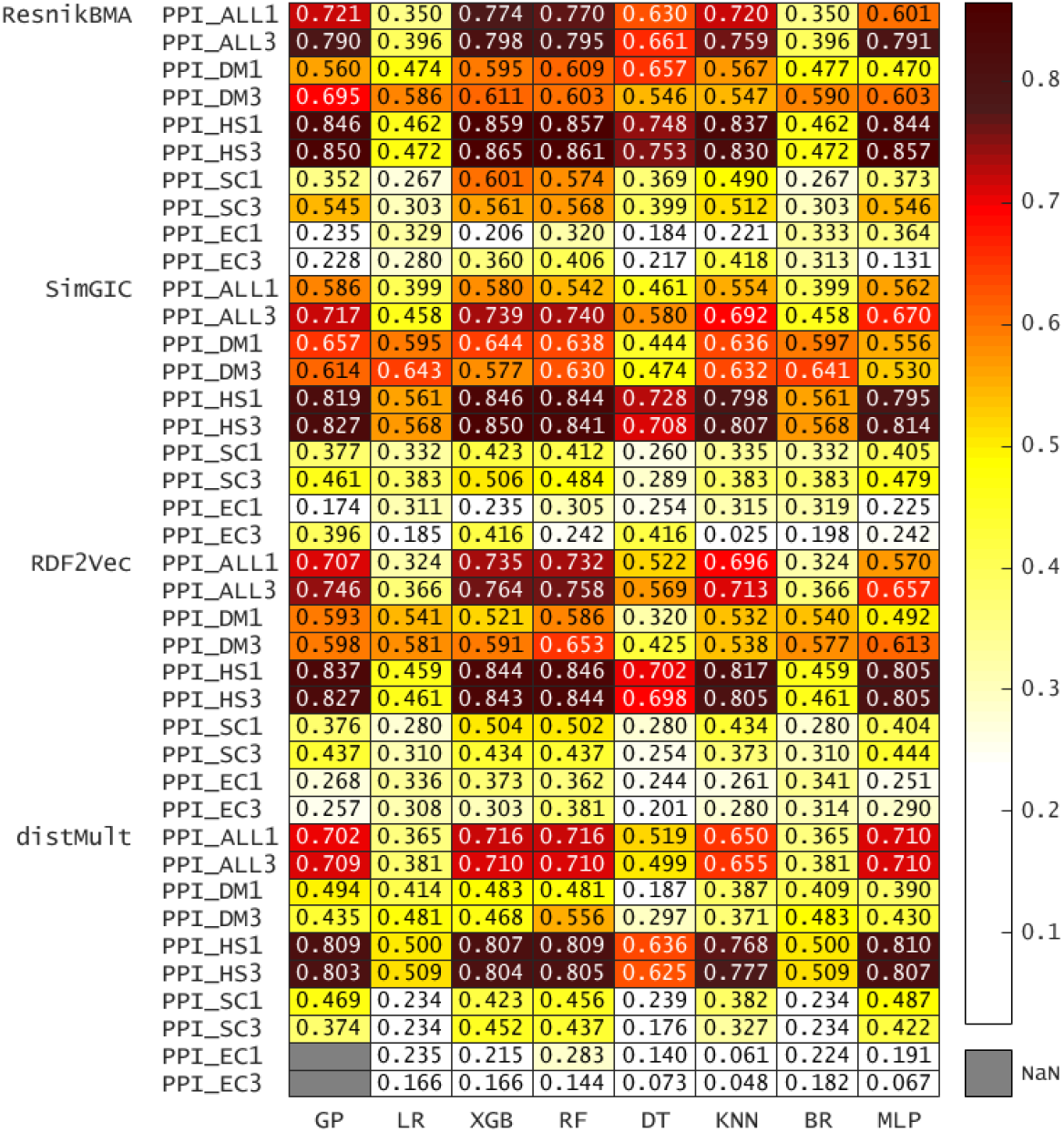
Heat map representing the median Pearson’s correlation coefficient using sequence proxy for PPI datasets.

**Figure 6.**
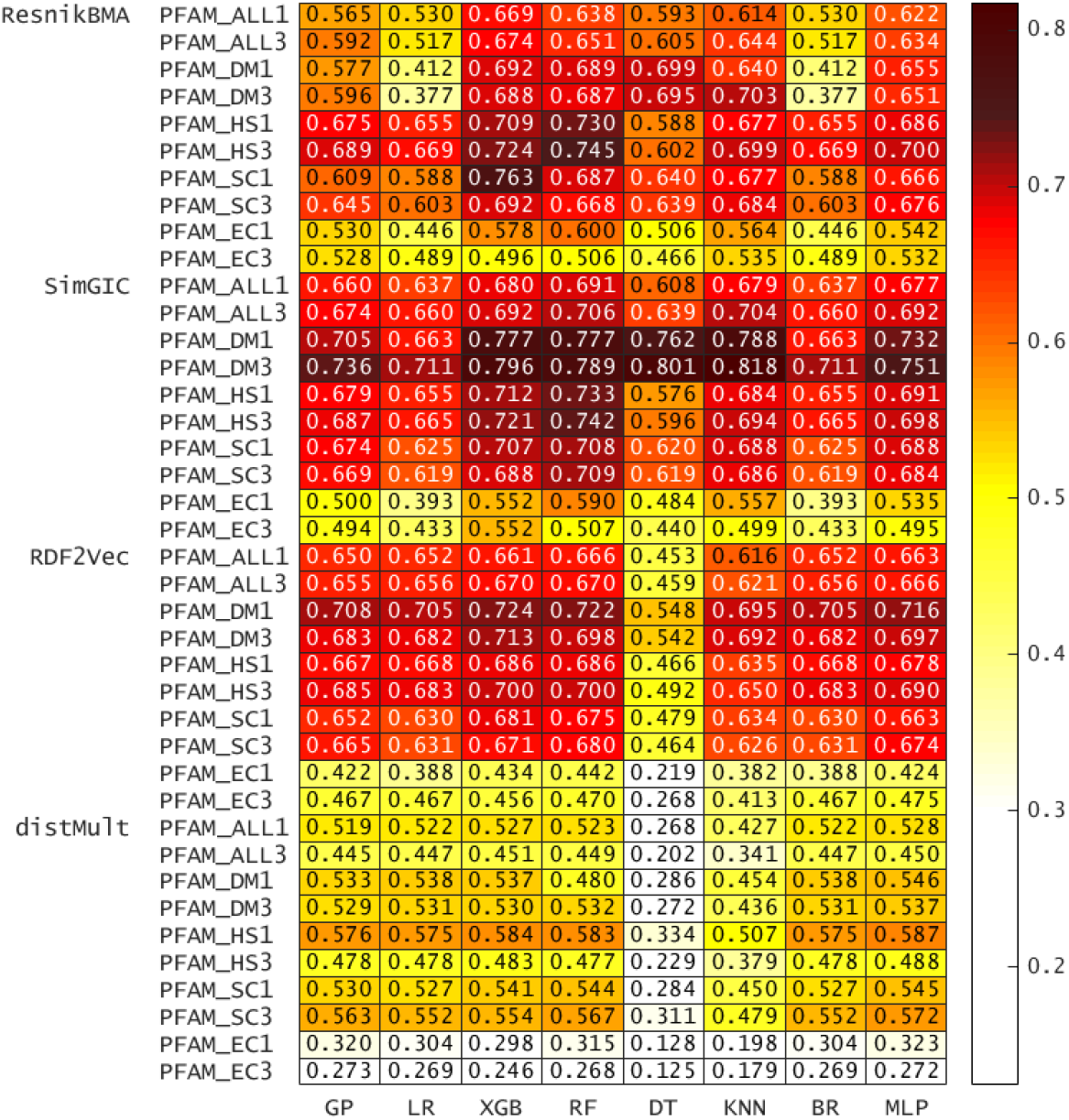
Heat map representing the median Pearson’s correlation coefficient using using PFAM proxy for PFAM datasets.

**Figure 7.**
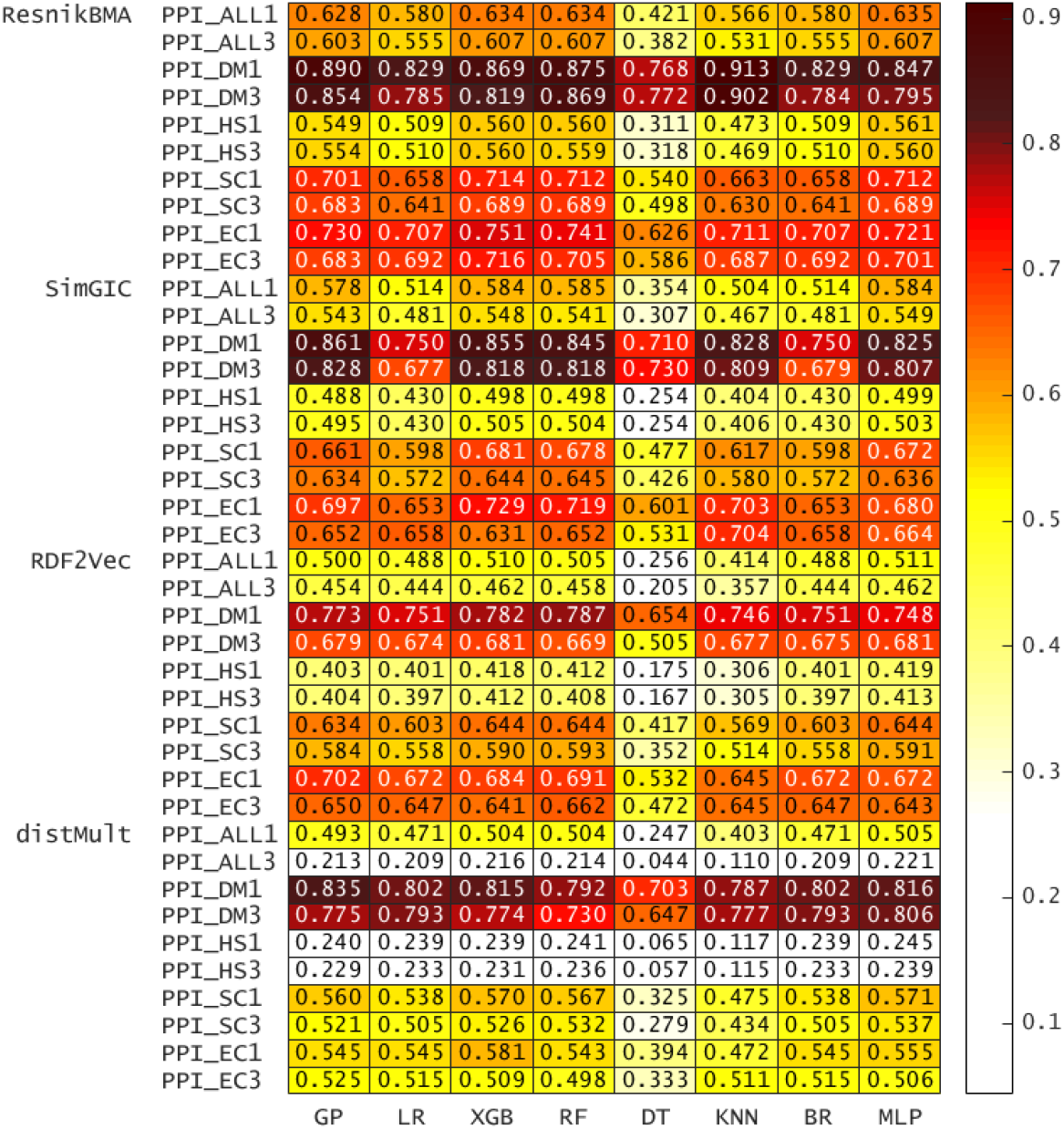
Heat map representing the median Pearson’s correlation coefficient using using PPI proxy for each PPI dataset.

**Figure 8.**
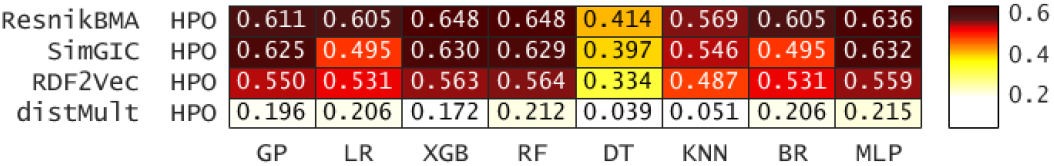
Heat map representing the median Pearson’s correlation coefficient using using phenotype series proxy for gene dataset.

**Figure 9.**
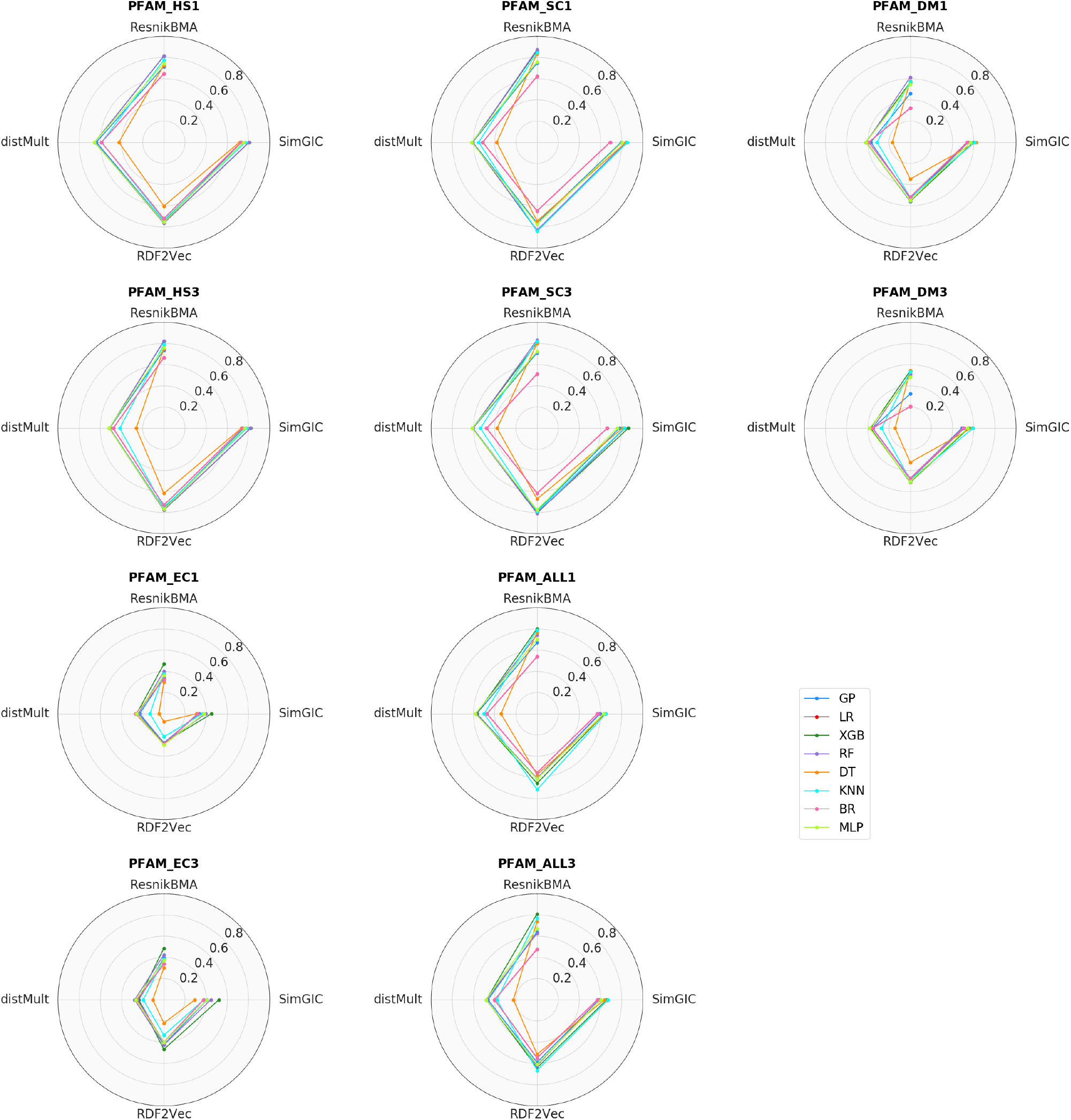
Radar charts using sequence proxy for PFAM datasets.

**Figure 10.**
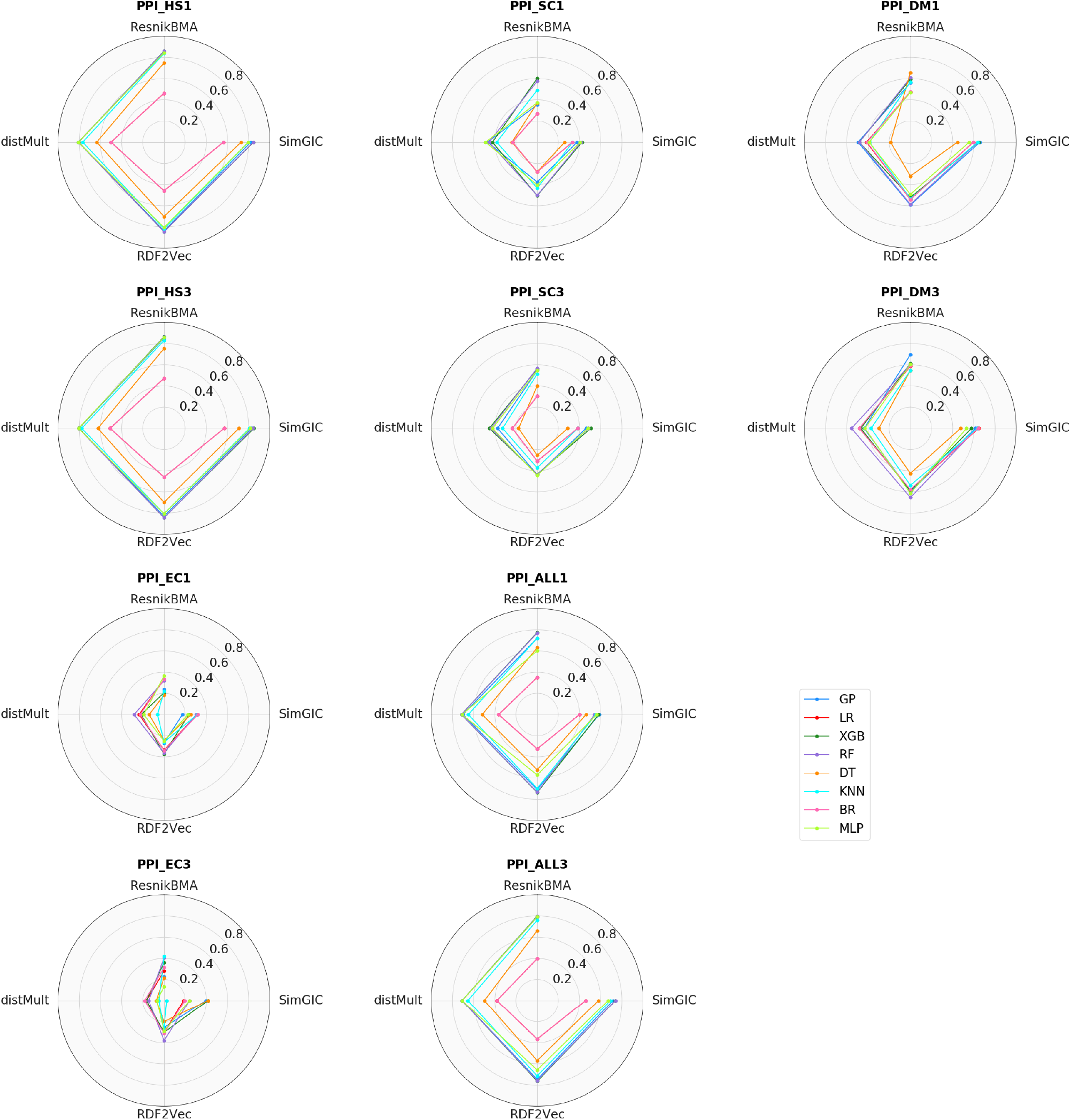
Radar charts using sequence proxy for PPI datasets.

**Figure 11.**
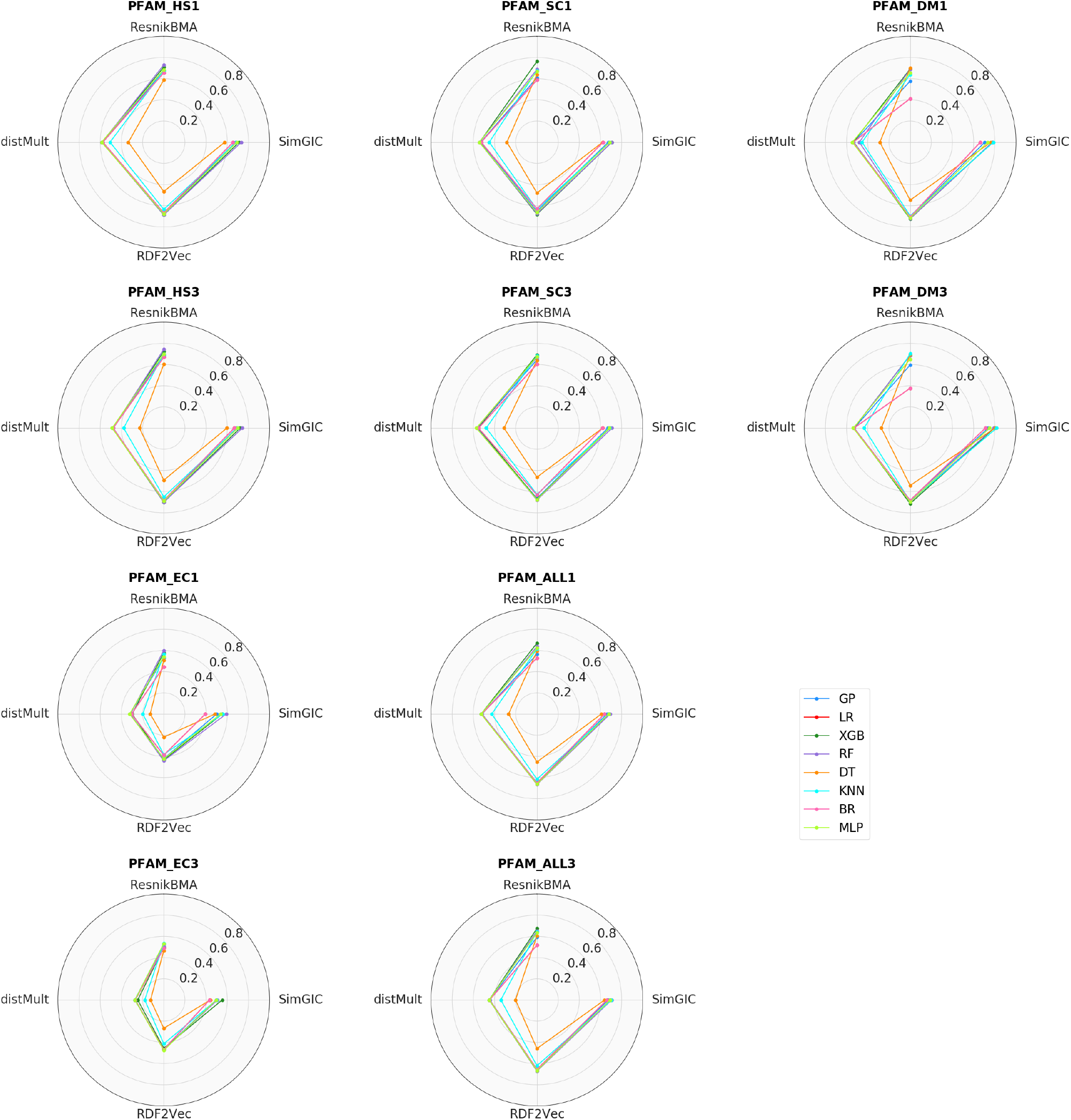
Radar charts using PFAM proxy for PFAM datasets.

**Figure 12.**
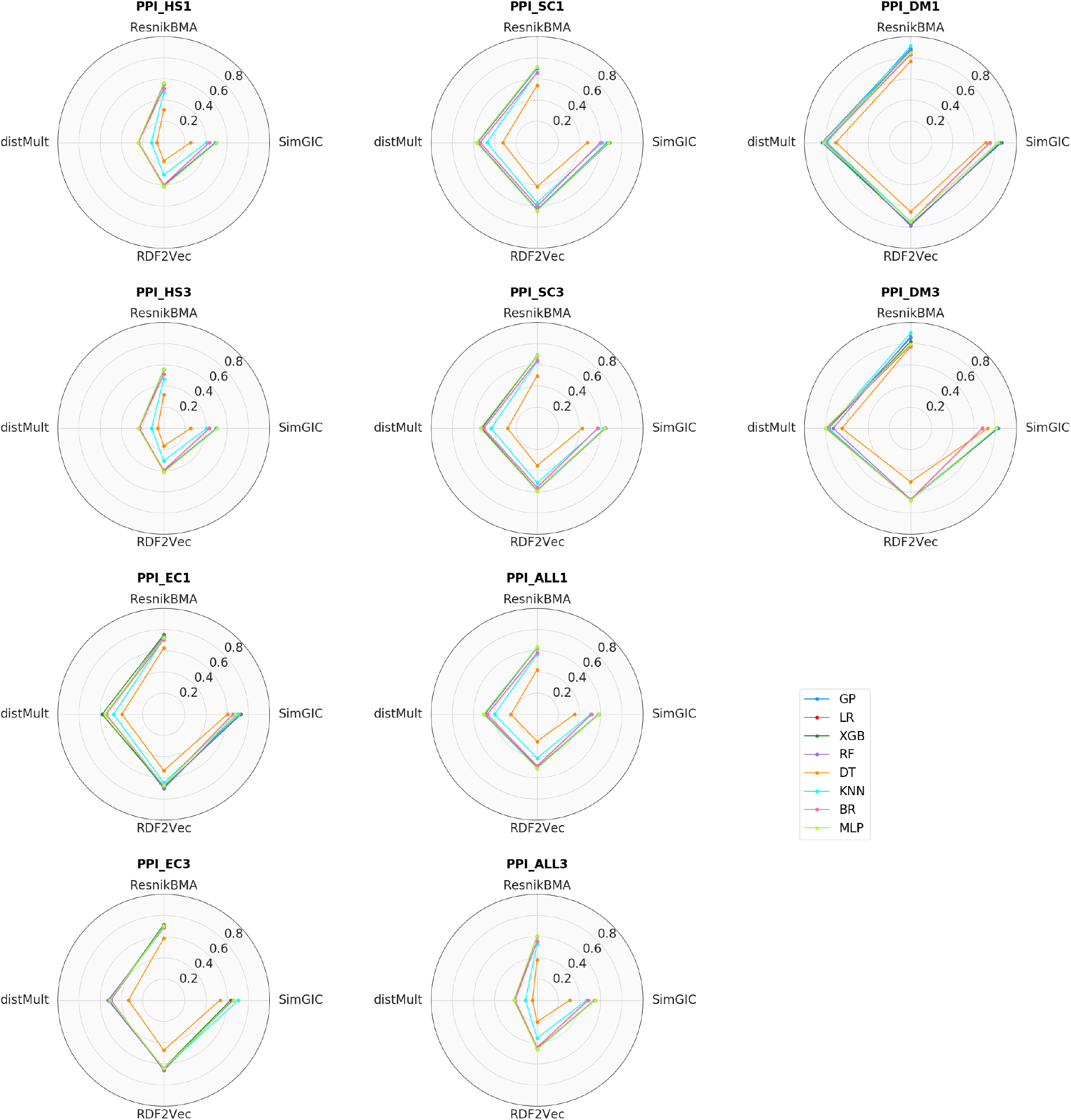
Radar charts using PPI proxy for PPI datasets.

**Figure 13.**
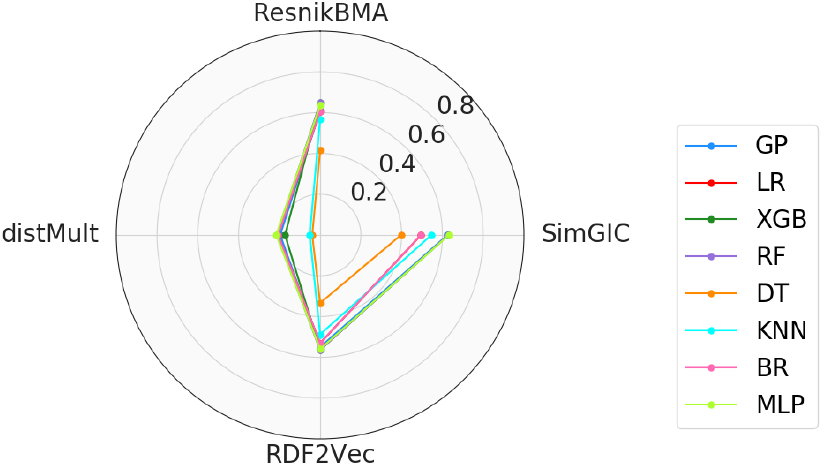
Radar charts using phenotype series proxy for gene dataset.

The performance of regression models obtained by DT is globally lower compared to the other ML algorithms. DT is one of the most commonly used approaches for supervised learning. However, since it is based on recursive binary splitting, DT may not be suitable for the current regression problem of finding the best combination of semantic aspects. LR and BR also show lower correlations in many cases. The Pearson correlation coefficients obtained by LR and BR are identical in most of the datasets. LR and BR assume a linear relationship between the independent and dependent variables, which is not valid for many cases. This characteristic may explain why these ML methods were not capable of learning suitable combinations of semantic aspects.

The very tight lines in the radar plots show that KNN, GP, and MLP achieve comparable results. Ensemble methods, like XGB and RF, achieve higher results in most experiments. These results are expected since the ensemble methods combine the decisions from multiple models to improve the overall performance, and these methods have been successfully applied to different domains [52].

Comparing the SSMs, the results seem to indicate that taxonomic semantic similarity is a more suitable similarity-based semantic representation for learning. Although the static similarity results have already demonstrated that taxonomic semantic similarity achieves higher correlations than graph embedding similarity, these differences are more evident when we apply ML methods.

A close investigation of the full results presented in Supplementary File also reveals interesting results regarding the significance tests. Statistically significant differences are determined using pairwise nonparametric Kruskal-Wallis tests at *p <* 0.01. Although MLP seems to achieve lower correlation values, there are no significant differences between XGB, RF or MLP and the best ML algorithm in most cases (which is either XGB or RF). Significant differences are more common between SSMs.

In order to assess whether a particular combination of an ML method and a specific SSM increases performance, for each proxy similarity we ranked the possible combinations of SSMs with ML algorithms within each dataset. Then, we calculated the average ranking of each SSM-ML combination. Table 5 shows the best combination for each proxy similarity, and they are always composed by a taxonomic SSM and an ensemble method.

**Table 5.** Best SSM-ML combination for each proxy similarity.

| Proxy | SSM | ML Algorithm |
| --- | --- | --- |
| Sim<sub>seq</sub> | ResnikBMA | RF |
| Sim<sub>PFAM</sub> | SimGIC | RF |
| PPI | ResnikBMA | XGB |
| Sim<sub>PS</sub> | ResnikBMA | XGB |

Choosing the right SSM and the right ML method for a particular application can be very hard. Several investigations have been handled to assess the performance of different SSMs, and different performances have been reported in different contexts [53]. Likewise, ML algorithms behave differently depending on many factors, from the type of problem at hand to the size of the data (No Free Lunch theorem [54]). Therefore, it is not straightforward to identify the best combination of SSM with ML algorithm that will work for all datasets and use cases. Nevertheless, the results in Table 5 seem to indicate that combining a taxonomic SSM with an ensemble method is a safe choice.

#### Supervised similarity interpretability

Although static SSMs, such as taxonomic SSMs, are hand-crafted and interpretable, supervised learning can lead to a loss of this valuable characteristic. Therefore, it is interesting to compare ML algorithms not only in terms of performance but also in terms of interpretability. The models obtained by KNN, BR, MLP and ensemble methods are more challenging to interpret, although some methods for explaining black-box models have been proposed [55]. In opposition, the LR models predict the target as a weighted sum of the feature inputs. These linear equations have an easy to understand interpretation. Table 6 shows, for each similarity proxy, a LR model obtained in one of the folds.

**Table 6.** Linear Regression models.

| Proxy | Model |
| --- | --- |
| Sim<sub>seq</sub> | 0.1450 SS<sub>BP</sub> + 0.0693 SS<sub>CC</sub> + 0.0916 SS<sub>MF</sub> - 0.0058 |
| Sim<sub>PFAM</sub> | 0.1943 SS<sub>BP</sub> + 0.1861 SS<sub>CC</sub> + 0.4344 SS<sub>MF</sub> + 0.1211 |
| PPI | 0.6001 SS<sub>BP</sub> + 0.6864 SS<sub>CC</sub> + 0.0132 SS<sub>MF</sub> + 0.1701 |
| Sim<sub>PS</sub> | 0.8406 SS<sub>HP</sub> + 0.2063 SS<sub>BP</sub> + 0.1282 SS<sub>CC</sub> + 0.0004 SS<sub>MF</sub> + 0.2261 |

The solutions obtained by DT and GP are also, in principle, interpretable. However, in both cases, trees may grow to be very complex while learning complicated datasets, which can raise some difficulty in interpreting the solutions. Fig 14 shows, for each similarity proxy, a GP model obtained in one of the folds. To allow a better understanding, these models were simplified to remove redundant and inviable code. Although the frequency in which a given variable appears in a GP model does not necessarily measure its importance for the predictions, the GP model analysis can indicate which semantic aspects are most relevant for each similarity proxy. The obtained DT models are very large with multiple levels deep, which decreases their interpretability and visualization and are thus not shown.

**Figure 14.**
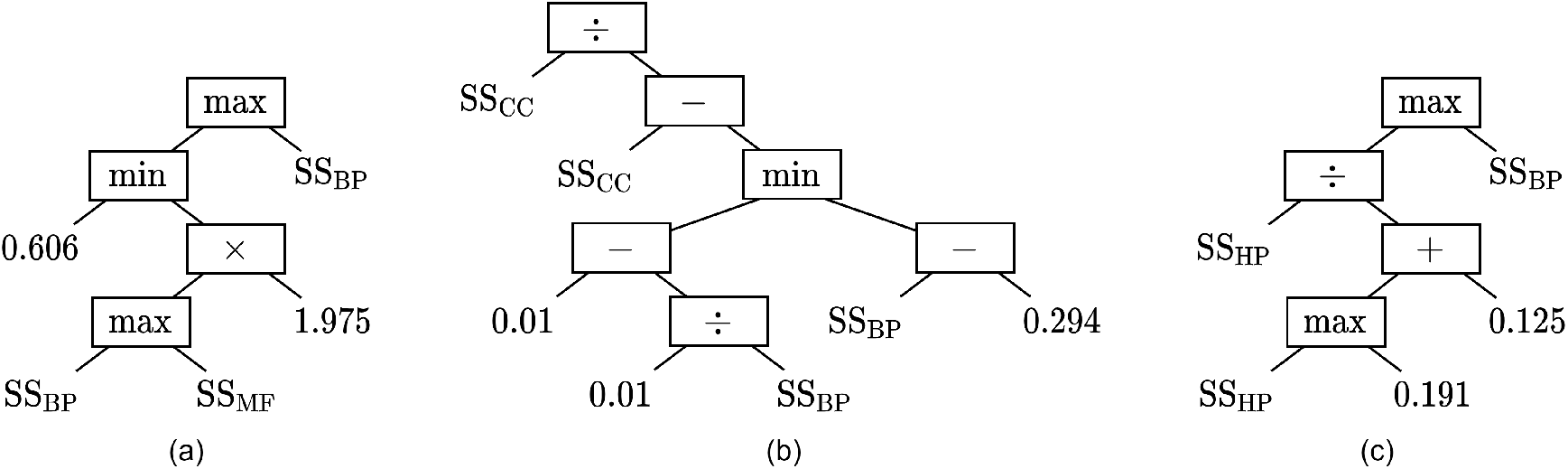
Parse trees representing GP models are shown for: (a) Sim_PFAM_; (b) PPI; (c) Sim_PS_.

It is important to note that, although interpretable models achieve lower performance values than blackbox models in most cases, as shown above, the supervised similarity obtained using LR and GP is still able to improve over the baselines.

### Static versus supervised similarity

Tables 7 to 10 compare the results obtained using static similarity and supervised similarity for sequence, PFAM, PPI and phenotypic series proxies, respectively. The static similarity was obtained using taxonomic SSMs (SimGIC or ResnikBMA), and then the Pearson correlation coefficient was computed for each proxy. Regarding supervised similarity, the median and IQR of Pearson correlation values were calculated for the proposed approach using a taxonomic SSM (SimGIC or ResnikBMA) with an ensemble method (XGB or RF) for each proxy, the combinations previously shown to produce the best results.

**Table 7.** Pearson correlation coefficient between Sim_seq_ and ResnikBMA or SimGIC for the baselines and the median and IQR of Pearson correlation coefficient between Sim_seq_ and supervised similarity obtained using XGB or RF. In bold, the best result for each dataset-SSM.

| Dataset | SSM | Static |  |  |  |  | Supervised |  |  |  |
| --- | --- | --- | --- | --- | --- | --- | --- | --- | --- | --- |
|  |  | BP | CC | MF | AVG | MAX | XGB |  | RF |  |
|  |  |  |  |  |  |  | Median | IQR | Median | IQR |
| PFAM |  |  |  |  |  |  |  |  |  |  |
| ALL1 | ResnikBMA | 0.528 | 0.373 | 0.291 | 0.481 | 0.399 | **0.803** | 0.013 | 0.746 | 0.015 |
|  | SimGIC | 0.552 | 0.406 | 0.415 | 0.547 | 0.406 | **0.640** | 0.033 | 0.589 | 0.004 |
| ALL3 | ResnikBMA | 0.466 | 0.334 | 0.325 | 0.445 | 0.349 | **0.810** | 0.012 | 0.626 | 0.009 |
|  | SimGIC | 0.544 | 0.374 | 0.451 | 0.539 | 0.411 | **0.658** | 0.037 | 0.580 | 0.009 |
| PPI |  |  |  |  |  |  |  |  |  |  |
| ALL1 | ResnikBMA | 0.258 | 0.222 | 0.326 | 0.323 | 0.250 | **0.774** | 0.024 | 0.770 | 0.028 |
|  | SimGIC | 0.317 | 0.257 | 0.370 | 0.380 | 0.280 | **0.580** | 0.030 | 0.542 | 0.029 |
| ALL3 | ResnikBMA | 0.293 | 0.260 | 0.370 | 0.376 | 0.289 | **0.798** | 0.044 | 0.795 | 0.039 |
|  | SimGIC | 0.371 | 0.295 | 0.411 | 0.439 | 0.316 | 0.739 | 0.031 | **0.740** | 0.025 |

These results show that whatever the ensemble method and taxonomic SSM, supervised similarity always achieves higher values of correlation than static similarity. Improvements over the single aspect similarities are consistent for all datasets and also clear when considering the combination baselines. However, there are some differences between the similarity proxies. For sequence proxy, it is known that the relationship between sequence similarity and semantic similarity is non-linear [56], so improvements over the best static similarity are very pronounced (up to 58% for PPI ALL1). Regarding PFAM proxy, supervised similarity outperforms both single aspects and static combinations (average and maximum), although the improvements are more relevant for single aspects. Concerning PPI proxy, improvements over the single aspect baselines are, as expected, more pronounced for the MF baseline (between 44 and 47%). In the gene dataset, the differences between static and supervised similarity are much more accentuated for the GO single aspects.

For the sake of brevity, Tables 7, 8, and 9 only show the results for the protein datasets combining all species’ protein pairs in the same group proxy.

**Table 8.** Pearson correlation coefficient between Sim_PFAM_ and ResnikBMA or SimGIC for the baselines and the median and IQR of Pearson correlation coefficient between Sim_PFAM_ and supervised similarity obtained using XGB or RF. In bold, the best result for each dataset-SSM.

| Dataset | SSM | Static |  |  |  |  | Supervised |  |  |  |
| --- | --- | --- | --- | --- | --- | --- | --- | --- | --- | --- |
|  |  | BP | CC | MF | AVG | MAX | XGB |  | RF |  |
|  |  |  |  |  |  |  | Median | IQR | Median | IQR |
| PFAM |  |  |  |  |  |  |  |  |  |  |
| ALL1 | ResnikBMA | 0.448 | 0.370 | 0.456 | 0.525 | 0.500 | **0.669** | 0.008 | 0.638 | 0.005 |
|  | SimGIC | 0.494 | 0.451 | 0.591 | 0.621 | 0.604 | 0.680 | 0.015 | **0.691** | 0.003 |
| ALL3 | ResnikBMA | 0.431 | 0.387 | 0.463 | 0.514 | 0.480 | **0.674** | 0.015 | 0.651 | 0.005 |
|  | SimGIC | 0.506 | 0.498 | 0.608 | 0.644 | 0.622 | 0.692 | 0.009 | **0.706** | 0.005 |

**Table 9.** Pearson correlation coefficient between PPI and ResnikBMA or SimGIC for the baselines and the median and IQR of Pearson correlation coefficient between PPI and supervised similarity obtained using XGB or RF. In bold, the best result for each dataset-SSM.

| Dataset | SSM | Static |  |  |  |  | Supervised |  |  |  |
| --- | --- | --- | --- | --- | --- | --- | --- | --- | --- | --- |
|  |  | BP | CC | MF | AVG | MAX | XGB |  | RF |  |
|  |  |  |  |  |  |  | Median | IQR | Median | IQR |
| PPI |  |  |  |  |  |  |  |  |  |  |
| ALL1 | ResnikBMA | 0.545 | 0.486 | 0.353 | 0.569 | 0.545 | **0.634** | 0.015 | 0.634 | 0.015 |
|  | SimGIC | 0.486 | 0.428 | 0.318 | 0.496 | 0.458 | 0.584 | 0.008 | **0.585** | 0.007 |
| ALL3 | ResnikBMA | 0.523 | 0.446 | 0.330 | 0.545 | 0.507 | **0.607** | 0.017 | 0.607 | 0.015 |
|  | SimGIC | 0.457 | 0.390 | 0.290 | 0.462 | 0.416 | **0.548** | 0.011 | 0.541 | 0.013 |

**Table 10.**
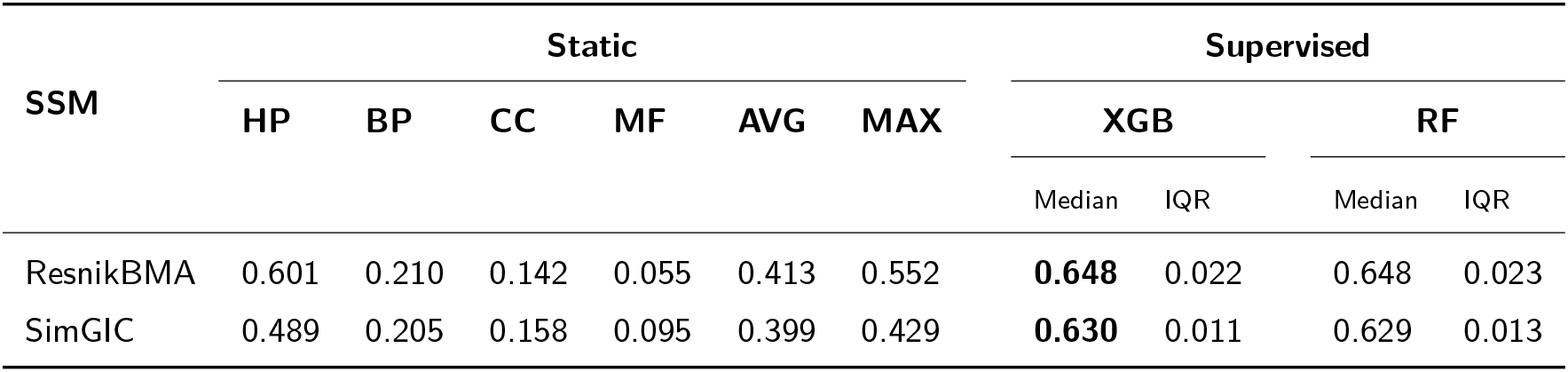
Pearson correlation coefficient between Sim_PS_ and ResnikBMA or SimGIC for the baselines and the median and IQR of Pearson correlation coefficient between Sim_PS_ and supervised similarity obtained using XGB or RF. In bold, the best result for each dataset-SSM.

However, Tables S4, S5 and S6 of Supplementary File provide the results for the remaining protein datasets and shows that supervised similarity, obtained with an ensemble method coupled with a taxonomic SSM, achieves better results than static similarity. Furthermore, we verify that, also for embedding similarity, our approach can learn a combination of semantic aspects that outperforms the best static similarity.

Finally, the comparison of results using protein datasets with different levels of annotation completion can be interesting. It is known that the biological entities should be well characterized in the context of the ontology to have enough information to compute semantic similarity between them. We have used benchmark datasets with two levels of annotation completion: the datasets ending in “1” include proteins with leaf-class annotations in, at least, one aspect; the datasets ending in “3” include proteins with at least one leaf-class annotation in each aspect. Analyzing our results, we conclude that in the PFAM datasets, lower correlations were generally found for the incomplete annotation datasets, but the opposite happens in the PPI datasets. These results are in agreement with conclusions in [57].

## Conclusion

Measuring the similarity between two gene products is a fundamental aspect of today’s biomedical informatics research. Biomedical ontologies and KGs provide meaningful context to data and support the comparison of biomedical entities through semantic similarity. Many KGs afford different perspectives over the data, however, existing SSMs are general-purpose and typically depend on expert knowledge to select and combine the relevant KG semantic aspects for each use case. Tailoring semantic similarity to a viewpoint of the domain or a particular use case in an automated fashion had not yet been tackled.

We have developed an approach that considers the different KG semantic aspects used to describe entities and relies on ML to learn a supervised semantic similarity. Currently, our approach includes five KG-based similarity measures based on embeddings or taxonomic semantic similarity, and eight ML methods. A comparative evaluation of the five SSMs combined with the eight ML algorithms was conducted using 21 benchmark datasets divided by species, level of annotation completion, KGs describing them, and similarity proxies employed in them. The similarity proxies rely on biological similarity measures and features - protein-protein interaction, protein function similarity, protein sequence similarity and phenotype-based gene similarity - and were used to train and test the supervised models. This evaluation elucidated which combinations of SSMs with ML algorithms are more suitable for each biological similarity.

The results showed that our approach is able to learn a supervised semantic similarity that outperforms static semantic similarity both using KG embeddings and standard taxonomic SSMs, obtaining more accurate similarity values. We specifically highlight the ability of the proposed approach to find better semantic aspect combination functions than static combinations emulating expert knowledge.

Our approach is independent of the SSM and the chosen ML method. Until now, we have used SSMs that take into consideration semantic and structural information. Recently, embedding methods, such as OPA2Vec [21], that also consider lexical information can be implemented and incorporated into our methodology. However, our goal is to select appropriate semantic aspect combinations to compute similarity, meaning that it is likely our approach not benefit so much from OPA2Vec.

Since interpretability has become a significant concern in ML, this issue should also be addressed in the future. Although we have employed interpretable ML algorithms, black-box models usually produced predictions with higher accuracy in our experiments, highlighting the need to explore the trade-off between performance and interpretability.

Although in this work we applied supervised ML algorithms to tailor semantic similarity to different similarity proxies, the proposed approach is versatile and can also be applied to tailor semantic similarity to a specific learning task. Consequently, there are multiple real-world tasks, where KG-based similarity is a suitable instance representation, that can benefit from this work. In future work, the impact of supervised similarity in tasks such as predicting protein-protein interactions, drug-target interactions or gene-disease associations should be evaluated.

## Acknowledgements

Not applicable.

## Funding

CP, SS, RTS are funded by the FCT through LASIGE Research Unit, ref. UIDB/00408/2020 and ref. UIDP/00408/2020. CP and RTS are funded by project SMILAX (ref. PTDC/EEI-ESS/4633/2014), SS by projects BINDER (ref. PTDC/CCI-INF/29168/2017) and PREDICT (ref. PTDC/CCI-CIF/29877/2017), and RTS by FCT PhD grant (ref. SFRH/BD/145377/2019). It was also partially supported by the KATY project which has received funding from the European Union’s Horizon 2020 research and innovation programme under grant agreement No 101017453.

### Abbreviations

BP: Biological Process
BR: Bayesian Ridge
CC: Cellular Component
DT: Decision Tree
GO: Gene Ontology
GP: Genetic Programming
HP: Human Phenotype Ontology
KG: Knowledge Graph
IC: Information Content
QR: Interquartile Range
KNN: *K*-Nearest Neighbor
LR: LinearRegression
MF: Molecular Function
ML: Machine Learning
MLP: Multi-Layer Perception
PPI: Protein-Protein Interaction
RF: Random Forest
SSM: Semantic Similarity Measure
XGB: XGBoost.

## Availability of data and materials

The datasets analysed during the current study are available in the GitHub repository (https://github.com/liseda-lab/kgsim-benchmark). The source code is also available in the GitHub repository (https://github.com/liseda-lab/Supervised-SS).

## Consent for publication

Not applicable.

## Authors’ contributions

All authors read and approved the final manuscript.

## Author details

LASIGE, Faculdade de Ciências da Universidade de Lisboa, Portugal.

## Additional Files

S1 File.

Supplementary tables.

## Footnotes

[1] https://github.com/liseda-lab/Supervised-SS

[2] https://github.com/liseda-lab/kgsim-benchmark

[3] https://github.com/IBCNServices/pyRDF2Vec

[4] https://github.com/thunlp/OpenKE/tree/OpenKE-Tensorflow1.0

[5] parameters for each embedding method are supplied in Supplementary File

[6] https://gplearn.readthedocs.io/en/stable/

[7] https://xgboost.readthedocs.io

[8] optimized parameters are supplied in Supplementary File

[9] In most of the radar charts, the red line representing LR is not visible due to the overlapping of the BR pink line.

