## Supplementary File for "Supervised Semantic Similarity"

RESEARCH

### Supervised Semantic Similarity - Supplementary Material

#### Parameters for knowledge graph embeddings

The size of each protein vector for each aspect was set to 200 for all methods. For RDF2Vec, we built a set of sequences generated from Weisfeiler-Lehman subtree kernels. For the Weisfeiler-Lehman algorithm, we used walks with depth 4, and we extracted a limited number of 500 random walks for each protein. The corpora of sequences were used to build a Skip-Gram model with the default parameters. These parameters were selected by their improved performance on a set of diverse benchmark data.For the remaining approaches, the default parameters given by OpenKE were used for TransE, TransH, TransD, TransR, distMult, and ComplEx models.

#### Parameters for ML algorithms

Table S1: Genetic Programming parameters.

| Parameter | Value |
|---|---|
| Population size | 500 |
| Number of generations | 50 |
| Fitness Function | RMSE |
| Tournament size | 20 |
| Stopping criteria | 0.0 |
| Range of constants to include in formulas | [-1.0,1.0] |
| Range of tree depths for the initial population | [2,6] |
| Initialization method | half and half |
| Probability of crossover on a tournament winner | 0.9 |
| Probability of subtree mutation on a tournament winner | 0.01 |
| Probability of hoist mutation on a tournament winner | 0.01 |
| Probability of point mutation on a tournament winner | 0.01 |
| Probability of any given node will be mutated, for point mutation | 0.05 |

Table S2: XGBoost parameters that have been optimized.

| Parameter | Values |
|---|---|
| Maximum depth | 2,4,6 |
| Number of estimators | 50,100,200 |
| Learning rate | 0.1, 0.01, 0.001 |

Table S3: Random Forest parameters that have been optimized.

| Parameters | Value |
|---|---|
| Maximum depth | 2,4,6, None |
| Number of estimators | 50,100,200 |

#### Correlation Results

Tables S4, S5, S6 and S7 provide the median Pearson correlation coefficient between the similarity proxies and supervised similarity obtained with the eight ML methods. In some cases, the results obtained with GP are not shown because the GP model only contains constants and, therefore, the Pearson correlation coefficient is undefined (division by zero).

Table S4: Pearson correlation coefficient between $\text{Sim}_{\text{seq}}$ and static similarity for the baselines and the median Pearson correlation coefficient between$\text{Sim}_{\text{seq}}$ and supervised similarity. Within each dataset, for each ML algorithm, the best SSM is in bold. The SSM is underlined when, using the same ML algorithm, there are no significant differences between the SSM and the best SSM (using $\alpha = 0.01$). Within each dataset, for each SSM, gray shading indicates the best ML algorithm. The light gray indicates that, using the same SSM, there are no significant differences between the ML algorithm and the best ML algorithm (using $\alpha = 0.01$).

| | | Static Similarity | | | | | Supervised Similarity | | | | | | | |
|---|---|---|---|---|---|---|---|---|---|---|---|---|---|---|
| | | BP | CC | MF | AVG | MAX | GP | LR | XGB | RF | DT | KNN | BR | MLP |
| PFAM_ALL1 | ResnikBMA | 0.528 | 0.373 | 0.291 | 0.481 | 0.399 | **0.672** | 0.539 | **0.803** | **0.746** | **0.768** | **0.790** | 0.539 | **0.703** |
| | SimGIC | 0.552 | 0.406 | 0.415 | 0.547 | 0.406 | 0.596 | **0.570** | 0.640 | 0.589 | 0.651 | 0.650 | **0.570** | 0.629 |
| | RDF2Vec | 0.540 | 0.437 | 0.419 | 0.544 | 0.457 | 0.610 | 0.558 | 0.657 | 0.610 | 0.575 | 0.716 | 0.558 | 0.615 |
| | TransE | -0.020 | -0.017 | -0.001 | -0.021 | -0.011 | | 0.025 | 0.025 | 0.022 | 0.002 | 0.003 | 0.025 | 0.025 |
| | distMult | 0.398 | 0.236 | 0.322 | 0.467 | 0.429 | 0.570 | 0.473 | 0.570 | 0.577 | 0.341 | 0.501 | 0.473 | 0.583 |

Table S4 Continued from previous page

| | | Static Similarity | | | | | Supervised Similarity | | | | | | | |
|---|---|---|---|---|---|---|---|---|---|---|---|---|---|---|
| | | **BP** | **CC** | **MF** | **AVG** | **MAX** | **GP** | **LR** | **XGB** | **RF** | **DT** | **KNN** | **BR** | **MLP** |
| PFAM_ALL3 | ResnikBMA | 0.466 | 0.334 | 0.325 | 0.445 | 0.349 | **0.641** | 0.478 | **0.810** | **0.626** | **0.740** | **0.771** | 0.478 | **0.672** |
| | SimGIC | 0.544 | 0.374 | 0.451 | 0.539 | 0.411 | 0.586 | **0.569** | 0.658 | 0.580 | 0.638 | 0.671 | **0.569** | 0.603 |
| | RDF2Vec | 0.520 | 0.394 | 0.469 | 0.533 | 0.442 | 0.589 | 0.548 | 0.641 | 0.620 | 0.513 | 0.664 | 0.548 | 0.608 |
| | TransE | -0.010 | -0.022 | 0.010 | -0.012 | -0.002 | | 0.023 | 0.024 | 0.024 | 0.000 | 0.003 | 0.023 | 0.024 |
| | distMult | 0.382 | 0.184 | 0.011 | 0.341 | 0.380 | 0.464 | 0.400 | 0.478 | 0.478 | 0.224 | 0.379 | 0.400 | 0.482 |
| PFAM_DM1 | ResnikBMA | 0.244 | 0.223 | 0.297 | 0.316 | 0.316 | 0.459 | 0.322 | 0.566 | **0.614** | 0.576 | 0.572 | 0.322 | 0.545 |
| | SimGIC | 0.436 | 0.305 | 0.484 | 0.505 | 0.438 | **0.548** | **0.537** | **0.597** | 0.546 | **0.624** | **0.609** | **0.537** | **0.571** |
| | RDF2Vec | 0.443 | 0.339 | 0.485 | 0.499 | 0.455 | 0.531 | 0.518 | 0.557 | 0.556 | 0.347 | 0.520 | 0.518 | 0.551 |
| | TransE | 0.015 | 0.015 | -0.005 | 0.013 | 0.011 | | 0.025 | 0.005 | 0.022 | 0.003 | 0.006 | 0.025 | 0.025 |
| | distMult | 0.319 | 0.223 | 0.266 | 0.388 | 0.347 | 0.373 | 0.392 | 0.418 | 0.415 | 0.169 | 0.314 | 0.392 | 0.418 |
| PFAM_DM3 | ResnikBMA | 0.147 | 0.103 | 0.197 | 0.179 | 0.178 | 0.325 | 0.205 | 0.543 | **0.510** | 0.545 | 0.535 | 0.205 | 0.483 |
| | SimGIC | 0.382 | 0.279 | 0.472 | 0.455 | 0.405 | **0.487** | **0.498** | **0.552** | 0.509 | **0.577** | **0.593** | **0.498** | **0.541** |
| | RDF2Vec | 0.370 | 0.304 | 0.458 | 0.442 | 0.408 | 0.473 | 0.473 | 0.511 | 0.484 | 0.322 | 0.487 | 0.473 | 0.505 |
| | TransE | -0.005 | 0.008 | 0.008 | 0.006 | 0.008 | | 0.008 | 0.002 | 0.005 | -0.003 | -0.004 | 0.008 | 0.000 |
| | distMult | 0.279 | 0.217 | 0.253 | 0.358 | 0.326 | 0.354 | 0.356 | 0.374 | 0.356 | 0.142 | 0.271 | 0.356 | 0.384 |
| PFAM_HS1 | ResnikBMA | 0.608 | 0.490 | 0.497 | 0.637 | 0.546 | 0.715 | 0.647 | **0.812** | **0.814** | **0.726** | **0.777** | 0.647 | 0.744 |
| | SimGIC | 0.676 | 0.607 | 0.619 | 0.724 | 0.625 | 0.732 | **0.729** | 0.806 | 0.804 | 0.714 | 0.768 | **0.729** | 0.751 |
| | RDF2Vec | 0.682 | 0.602 | 0.626 | 0.717 | 0.645 | **0.751** | 0.721 | 0.763 | 0.762 | 0.603 | 0.737 | 0.721 | **0.756** |
| | TransE | -0.110 | -0.105 | -0.081 | -0.155 | -0.124 | | 0.151 | 0.151 | 0.151 | 0.025 | 0.055 | 0.151 | 0.155 |
| | distMult | 0.515 | 0.289 | 0.424 | 0.581 | 0.533 | 0.638 | 0.589 | 0.653 | 0.652 | 0.425 | 0.588 | 0.589 | 0.656 |
| PFAM_HS3 | ResnikBMA | 0.620 | 0.523 | 0.506 | 0.653 | 0.560 | 0.738 | 0.666 | **0.821** | **0.825** | **0.749** | **0.792** | 0.666 | 0.763 |
| | SimGIC | 0.690 | 0.623 | 0.629 | 0.734 | 0.633 | 0.754 | **0.740** | 0.820 | 0.817 | 0.732 | 0.783 | **0.740** | 0.767 |
| | RDF2Vec | 0.693 | 0.622 | 0.633 | 0.726 | 0.652 | **0.761** | 0.729 | 0.777 | 0.776 | 0.615 | 0.750 | 0.729 | **0.769** |
| | TransE | -0.126 | -0.117 | -0.086 | -0.171 | -0.132 | 0.158 | 0.174 | 0.176 | 0.173 | 0.028 | 0.078 | 0.174 | 0.181 |
| | distMult | 0.073 | 0.303 | 0.425 | 0.439 | 0.379 | 0.512 | 0.479 | 0.514 | 0.513 | 0.262 | 0.416 | 0.479 | 0.517 |
| PFAM_SC1 | ResnikBMA | 0.609 | 0.478 | 0.274 | 0.554 | 0.473 | 0.746 | 0.622 | **0.865** | **0.875** | 0.835 | 0.847 | 0.622 | 0.761 |
| | SimGIC | 0.675 | 0.522 | 0.369 | 0.616 | 0.398 | **0.794** | **0.690** | 0.841 | 0.841 | **0.840** | **0.855** | **0.690** | **0.804** |
| | RDF2Vec | 0.620 | 0.530 | 0.387 | 0.617 | 0.493 | 0.762 | 0.649 | 0.835 | 0.830 | 0.744 | 0.842 | 0.649 | 0.775 |
| | TransE | 0.008 | 0.006 | 0.029 | 0.024 | 0.021 | | 0.030 | 0.040 | 0.033 | 0.013 | 0.013 | 0.030 | 0.040 |
| | distMult | 0.417 | 0.337 | 0.347 | 0.511 | 0.449 | 0.608 | 0.513 | 0.609 | 0.609 | 0.380 | 0.551 | 0.513 | 0.617 |
| PFAM_SC3 | ResnikBMA | 0.497 | 0.378 | 0.333 | 0.480 | 0.357 | 0.712 | 0.512 | 0.808 | **0.833** | 0.800 | 0.822 | 0.512 | 0.725 |
| | SimGIC | 0.660 | 0.443 | 0.394 | 0.594 | 0.390 | 0.777 | **0.663** | **0.861** | 0.830 | **0.807** | **0.824** | **0.663** | 0.756 |
| | RDF2Vec | 0.597 | 0.449 | 0.443 | 0.591 | 0.454 | **0.782** | 0.613 | 0.801 | 0.809 | 0.668 | 0.790 | 0.613 | **0.775** |
| | TransE | 0.062 | 0.022 | 0.082 | 0.091 | 0.071 | 0.094 | 0.106 | 0.100 | 0.105 | 0.006 | 0.031 | 0.106 | 0.112 |
| | distMult | 0.387 | 0.247 | 0.377 | 0.475 | 0.440 | 0.606 | 0.476 | 0.610 | 0.607 | 0.377 | 0.536 | 0.476 | 0.614 |

Table S4 Continued from previous page

| | | Static Similarity | | | | | Supervised Similarity | | | | | | | |
|---|---|---|---|---|---|---|---|---|---|---|---|---|---|---|
| | | **BP** | **CC** | **MF** | **AVG** | **MAX** | **GP** | **LR** | **XGB** | **RF** | **DT** | **KNN** | **BR** | **MLP** |
| PFAM_EC1 | ResnikBMA | 0.276 | 0.168 | 0.309 | 0.313 | 0.254 | **0.344** | **0.325** | **0.468** | **0.400** | 0.300 | **0.378** | **0.325** | 0.361 |
| | SimGIC | 0.266 | 0.179 | 0.306 | 0.320 | 0.258 | 0.334 | 0.319 | 0.451 | 0.397 | **0.309** | 0.365 | 0.319 | **0.380** |
| | RDF2Vec | 0.237 | 0.167 | 0.234 | 0.284 | 0.210 | 0.275 | 0.288 | 0.273 | 0.276 | 0.074 | 0.217 | 0.288 | 0.294 |
| | TransE | 0.020 | 0.021 | 0.007 | 0.028 | 0.025 | | 0.021 | -0.019 | 0.020 | -0.031 | -0.004 | 0.021 | 0.025 |
| | distMult | 0.066 | 0.100 | 0.240 | 0.217 | 0.197 | 0.229 | 0.265 | 0.257 | 0.240 | 0.046 | 0.132 | 0.265 | 0.256 |
| PFAM_EC3 | ResnikBMA | 0.327 | 0.200 | 0.287 | 0.347 | 0.281 | 0.394 | 0.338 | 0.486 | 0.425 | **0.302** | **0.382** | 0.339 | 0.372 |
| | SimGIC | 0.368 | 0.219 | 0.304 | 0.392 | 0.297 | 0.410 | 0.375 | **0.520** | **0.443** | 0.293 | 0.378 | 0.374 | **0.409** |
| | RDF2Vec | 0.326 | 0.233 | 0.330 | 0.411 | 0.297 | **0.421** | **0.399** | 0.467 | 0.424 | 0.220 | 0.333 | **0.399** | 0.400 |
| | TransE | 0.014 | 0.028 | 0.023 | 0.037 | 0.016 | | 0.041 | -0.005 | -0.038 | 0.018 | -0.022 | 0.040 | 0.026 |
| | distMult | 0.129 | 0.140 | 0.237 | 0.275 | 0.250 | 0.237 | 0.253 | 0.240 | 0.276 | 0.104 | 0.194 | 0.253 | 0.265 |
| PPI_ALL1 | ResnikBMA | 0.258 | 0.222 | 0.326 | 0.323 | 0.250 | **0.721** | 0.350 | **0.774** | **0.770** | **0.630** | **0.720** | 0.350 | 0.601 |
| | SimGIC | 0.317 | 0.257 | 0.370 | 0.380 | 0.280 | 0.586 | 0.399 | 0.580 | 0.542 | 0.461 | 0.554 | 0.399 | 0.562 |
| | RDF2Vec | 0.274 | 0.237 | 0.297 | 0.316 | 0.268 | 0.707 | 0.324 | 0.735 | 0.732 | 0.522 | 0.696 | 0.324 | 0.570 |
| | TransE | 0.355 | 0.352 | 0.364 | 0.498 | 0.401 | 0.688 | **0.501** | 0.688 | 0.688 | 0.452 | 0.622 | **0.501** | 0.688 |
| | distMult | 0.277 | 0.239 | 0.202 | 0.369 | 0.310 | 0.702 | 0.365 | 0.716 | 0.716 | 0.519 | 0.650 | 0.365 | **0.710** |
| PPI_ALL3 | ResnikBMA | 0.293 | 0.260 | 0.370 | 0.376 | 0.289 | **0.790** | 0.396 | **0.798** | **0.795** | **0.661** | **0.759** | 0.396 | **0.791** |
| | SimGIC | 0.371 | 0.295 | 0.411 | 0.439 | 0.316 | 0.717 | 0.458 | 0.739 | 0.740 | 0.580 | 0.692 | 0.458 | 0.670 |
| | RDF2Vec | 0.304 | 0.266 | 0.330 | 0.358 | 0.300 | 0.746 | 0.366 | 0.764 | 0.758 | 0.569 | 0.713 | 0.366 | 0.657 |
| | TransE | 0.391 | 0.391 | 0.399 | 0.536 | 0.439 | 0.707 | **0.534** | 0.707 | 0.708 | 0.478 | 0.650 | **0.534** | 0.707 |
| | distMult | 0.228 | 0.262 | 0.225 | 0.375 | 0.285 | 0.709 | 0.381 | 0.710 | 0.710 | 0.499 | 0.655 | 0.381 | 0.710 |
| PPI_DM1 | ResnikBMA | 0.351 | 0.384 | 0.384 | 0.422 | 0.357 | 0.560 | 0.474 | 0.595 | 0.609 | **0.657** | 0.567 | 0.477 | 0.470 |
| | SimGIC | 0.393 | 0.449 | 0.526 | 0.514 | 0.451 | **0.657** | **0.595** | **0.644** | **0.638** | 0.444 | **0.636** | **0.597** | **0.556** |
| | RDF2Vec | 0.405 | 0.399 | 0.480 | 0.481 | 0.433 | 0.593 | 0.541 | 0.521 | 0.586 | 0.320 | 0.532 | 0.540 | 0.492 |
| | TransE | -0.054 | -0.033 | 0.012 | -0.040 | -0.004 | | -0.051 | -0.036 | -0.038 | -0.018 | 0.020 | -0.051 | -0.036 |
| | distMult | 0.244 | 0.287 | 0.323 | 0.358 | 0.307 | 0.494 | 0.414 | 0.483 | 0.481 | 0.187 | 0.387 | 0.409 | 0.390 |
| PPI_DM3 | ResnikBMA | 0.383 | 0.417 | 0.457 | 0.483 | 0.400 | **0.695** | 0.586 | **0.611** | 0.603 | **0.546** | 0.547 | 0.590 | 0.603 |
| | SimGIC | 0.454 | 0.461 | 0.571 | 0.556 | 0.487 | 0.614 | **0.643** | 0.577 | 0.630 | 0.474 | **0.632** | **0.641** | 0.530 |
| | RDF2Vec | 0.433 | 0.430 | 0.512 | 0.514 | 0.448 | 0.598 | 0.581 | 0.591 | **0.653** | 0.425 | 0.538 | 0.577 | **0.613** |
| | TransE | -0.086 | -0.044 | 0.013 | -0.064 | -0.008 | | -0.045 | -0.002 | -0.122 | 0.038 | 0.040 | -0.052 | -0.051 |
| | distMult | 0.250 | 0.272 | 0.347 | 0.373 | 0.311 | 0.435 | 0.481 | 0.468 | 0.556 | 0.297 | 0.371 | 0.483 | 0.430 |
| PPI_HS1 | ResnikBMA | 0.358 | 0.305 | 0.422 | 0.441 | 0.339 | **0.846** | 0.462 | **0.859** | **0.857** | **0.748** | **0.837** | 0.462 | **0.844** |
| | SimGIC | 0.484 | 0.371 | 0.489 | 0.538 | 0.390 | 0.819 | 0.561 | 0.846 | 0.844 | 0.728 | 0.798 | 0.561 | 0.795 |
| | RDF2Vec | 0.377 | 0.333 | 0.383 | 0.451 | 0.374 | 0.837 | 0.459 | 0.844 | 0.846 | 0.702 | 0.817 | 0.459 | 0.805 |
| | TransE | 0.492 | 0.497 | 0.484 | 0.640 | 0.535 | 0.805 | **0.639** | 0.805 | 0.805 | 0.645 | 0.770 | **0.639** | 0.805 |
| | distMult | 0.313 | 0.331 | 0.351 | 0.502 | 0.390 | 0.809 | 0.500 | 0.807 | 0.809 | 0.636 | 0.768 | 0.500 | 0.810 |

Table S4 Continued from previous page

| | | **Static Similarity** | | | | | **Supervised Similarity** | | | | | | | |
|---|---|---|---|---|---|---|---|---|---|---|---|---|---|---|
| | | **BP** | **CC** | **MF** | **AVG** | **MAX** | **GP** | **LR** | **XGB** | **RF** | **DT** | **KNN** | **BR** | **MLP** |
| PPI_HS3 | ResnikBMA | 0.367 | 0.313 | 0.432 | 0.452 | 0.347 | **0.850** | 0.472 | **0.865** | **0.861** | **0.753** | **0.830** | 0.472 | **0.857** |
| | SimGIC | 0.494 | 0.377 | 0.497 | 0.548 | 0.396 | 0.827 | 0.568 | 0.850 | 0.841 | 0.708 | 0.807 | 0.568 | 0.814 |
| | RDF2Vec | 0.382 | 0.337 | 0.395 | 0.456 | 0.382 | 0.827 | 0.461 | 0.843 | 0.844 | 0.698 | 0.805 | 0.461 | 0.805 |
| | TransE | 0.498 | 0.503 | 0.491 | 0.646 | 0.540 | 0.802 | **0.650** | 0.802 | 0.802 | 0.627 | 0.756 | **0.650** | 0.802 |
| | distMult | 0.319 | 0.337 | 0.357 | 0.509 | 0.397 | 0.803 | 0.509 | 0.804 | 0.805 | 0.625 | 0.777 | 0.509 | 0.807 |
| PPI_SC1 | ResnikBMA | 0.192 | 0.153 | 0.259 | 0.235 | 0.194 | 0.352 | 0.267 | **0.601** | **0.574** | **0.369** | **0.490** | 0.267 | 0.373 |
| | SimGIC | 0.254 | 0.188 | 0.307 | 0.303 | 0.229 | 0.377 | **0.332** | 0.423 | 0.412 | 0.260 | 0.335 | **0.332** | 0.405 |
| | RDF2Vec | 0.235 | 0.184 | 0.253 | 0.265 | 0.227 | 0.376 | 0.280 | 0.504 | 0.502 | 0.280 | 0.434 | 0.280 | 0.404 |
| | TransE | 0.077 | 0.075 | 0.087 | 0.127 | 0.105 | 0.294 | 0.136 | 0.296 | 0.307 | 0.087 | 0.170 | 0.136 | 0.308 |
| | distMult | 0.185 | 0.129 | 0.172 | 0.234 | 0.214 | **0.469** | 0.234 | 0.423 | 0.456 | 0.239 | 0.382 | 0.234 | **0.487** |
| PPI_SC3 | ResnikBMA | 0.211 | 0.179 | 0.293 | 0.273 | 0.220 | **0.545** | 0.303 | **0.561** | **0.568** | **0.399** | **0.512** | 0.303 | **0.546** |
| | SimGIC | 0.292 | 0.215 | 0.342 | 0.354 | 0.262 | 0.461 | **0.383** | 0.506 | 0.484 | 0.289 | 0.383 | **0.383** | 0.479 |
| | RDF2Vec | 0.257 | 0.207 | 0.278 | 0.301 | 0.253 | 0.437 | 0.310 | 0.434 | 0.437 | 0.254 | 0.373 | 0.310 | 0.444 |
| | TransE | 0.066 | 0.065 | 0.069 | 0.107 | 0.087 | 0.014 | 0.039 | 0.032 | 0.027 | -0.001 | -0.008 | 0.039 | 0.048 |
| | distMult | 0.196 | 0.137 | 0.176 | 0.248 | 0.227 | 0.374 | 0.234 | 0.452 | 0.437 | 0.176 | 0.327 | 0.234 | 0.422 |
| PPI_EC1 | ResnikBMA | 0.170 | 0.201 | 0.215 | 0.229 | 0.209 | 0.235 | 0.329 | 0.206 | 0.320 | 0.184 | 0.221 | 0.333 | **0.364** |
| | SimGIC | 0.173 | 0.164 | 0.236 | 0.225 | 0.235 | 0.174 | 0.311 | 0.235 | 0.305 | **0.254** | **0.315** | 0.319 | 0.225 |
| | RDF2Vec | 0.171 | 0.147 | 0.269 | 0.238 | 0.242 | **0.268** | **0.336** | **0.373** | **0.362** | 0.244 | 0.261 | **0.341** | 0.251 |
| | TransE | -0.048 | 0.012 | 0.024 | -0.007 | -0.011 | | -0.060 | -0.025 | -0.073 | -0.063 | 0.005 | -0.059 | -0.035 |
| | distMult | 0.091 | 0.136 | 0.088 | 0.152 | 0.131 | | 0.235 | 0.215 | 0.283 | 0.140 | 0.061 | 0.224 | 0.191 |
| PPI_EC3 | ResnikBMA | 0.104 | 0.144 | 0.180 | 0.168 | 0.145 | 0.228 | 0.280 | 0.360 | **0.406** | 0.217 | **0.418** | 0.313 | 0.131 |
| | SimGIC | 0.136 | 0.088 | 0.221 | 0.171 | 0.185 | **0.396** | 0.185 | **0.416** | 0.242 | **0.416** | 0.025 | 0.198 | 0.242 |
| | RDF2Vec | 0.123 | 0.064 | 0.247 | 0.176 | 0.199 | 0.257 | **0.308** | 0.303 | 0.381 | 0.201 | 0.280 | **0.314** | **0.290** |
| | TransE | -0.054 | 0.051 | 0.052 | 0.025 | 0.019 | | 0.049 | -0.079 | -0.107 | -0.016 | -0.085 | 0.047 | 0.017 |
| | distMult | 0.057 | 0.087 | 0.094 | 0.114 | 0.090 | | 0.166 | 0.166 | 0.144 | 0.073 | 0.048 | 0.182 | 0.067 |

Table S5: Pearson correlation coefficient between $\text{sim}_{\text{PFAM}}$ and static similarity for the baselines and the median Pearson correlation coefficient between $\text{sim}_{\text{PFAM}}$ and supervised similarity. Within each dataset, for each ML algorithm, the best SSM is in bold. The SSM is underlined when, using the same ML algorithm, there are no significant differences between the SSM and the best SSM (using $\alpha = 0.01$). Within each dataset, for each SSM, gray shading indicates the best ML algorithm. The light gray indicates that, using the same SSM, there are no significant differences between the ML algorithm and the best ML algorithm (using $\alpha = 0.01$).

| | | **Static Similarity** | | | | | **Supervised Similarity** | | | | | | | |
|---|---|---|---|---|---|---|---|---|---|---|---|---|---|---|
| | | **BP** | **CC** | **MF** | **AVG** | **MAX** | **GP** | **LR** | **XGB** | **RF** | **DT** | **KNN** | **BR** | **MLP** |
| PFAM_ALL1 | ResnikBMA | 0.448 | 0.370 | 0.456 | 0.525 | 0.500 | 0.565 | 0.530 | 0.669 | 0.638 | 0.593 | 0.614 | 0.530 | 0.622 |
| | SimGIC | 0.494 | 0.451 | 0.591 | 0.621 | 0.604 | **0.660** | 0.637 | **0.680** | **0.691** | **0.608** | **0.679** | 0.637 | **0.677** |
| | RDF2Vec | 0.524 | 0.466 | 0.619 | 0.627 | 0.623 | 0.650 | **0.652** | 0.661 | 0.666 | 0.453 | 0.616 | **0.652** | 0.663 |
| | TransE | -0.032 | -0.031 | -0.011 | -0.039 | -0.027 | | 0.042 | 0.039 | 0.036 | -0.001 | 0.006 | 0.042 | 0.042 |
| | distMult | 0.414 | 0.254 | 0.388 | 0.516 | 0.457 | 0.519 | 0.522 | 0.527 | 0.523 | 0.268 | 0.427 | 0.522 | 0.528 |
| PFAM_ALL3 | ResnikBMA | 0.431 | 0.387 | 0.463 | 0.514 | 0.480 | 0.592 | 0.517 | 0.674 | 0.651 | 0.605 | 0.644 | 0.517 | 0.634 |
| | SimGIC | 0.506 | 0.498 | 0.608 | 0.644 | 0.622 | **0.674** | **0.660** | **0.692** | **0.706** | **0.639** | **0.704** | **0.660** | **0.692** |
| | RDF2Vec | 0.535 | 0.514 | 0.612 | 0.640 | 0.627 | 0.655 | 0.656 | 0.670 | 0.670 | 0.459 | 0.621 | 0.656 | 0.666 |
| | TransE | -0.005 | -0.046 | -0.020 | -0.039 | -0.033 | | 0.048 | 0.044 | 0.044 | -0.003 | 0.006 | 0.048 | 0.047 |
| | distMult | 0.413 | 0.242 | 0.036 | 0.400 | 0.378 | 0.445 | 0.447 | 0.451 | 0.449 | 0.202 | 0.341 | 0.447 | 0.450 |
| PFAM_DM1 | ResnikBMA | 0.289 | 0.306 | 0.372 | 0.402 | 0.414 | 0.577 | 0.412 | 0.692 | 0.689 | 0.699 | 0.640 | 0.412 | 0.655 |
| | SimGIC | 0.495 | 0.453 | 0.600 | 0.644 | 0.617 | 0.705 | 0.663 | **0.777** | **0.777** | **0.762** | **0.788** | 0.663 | **0.732** |
| | RDF2Vec | 0.526 | 0.480 | 0.676 | 0.663 | 0.658 | **0.708** | **0.705** | 0.724 | 0.722 | 0.548 | 0.695 | **0.705** | 0.716 |
| | TransE | -0.025 | -0.007 | -0.013 | -0.025 | -0.020 | | 0.025 | 0.016 | 0.020 | 0.002 | -0.006 | 0.025 | 0.024 |
| | distMult | 0.390 | 0.279 | 0.438 | 0.531 | 0.475 | 0.533 | 0.538 | 0.537 | 0.480 | 0.286 | 0.454 | 0.538 | 0.546 |
| PFAM_DM3 | ResnikBMA | 0.207 | 0.302 | 0.343 | 0.345 | 0.330 | 0.596 | 0.377 | 0.688 | 0.687 | 0.695 | 0.703 | 0.377 | 0.651 |
| | SimGIC | 0.518 | 0.565 | 0.612 | 0.695 | 0.656 | **0.736** | **0.711** | **0.796** | **0.789** | **0.801** | **0.818** | **0.711** | **0.751** |
| | RDF2Vec | 0.510 | 0.560 | 0.608 | 0.661 | 0.646 | 0.683 | 0.682 | 0.713 | 0.698 | 0.542 | 0.692 | 0.682 | 0.697 |
| | TransE | -0.002 | -0.027 | -0.006 | -0.020 | -0.019 | | 0.027 | 0.019 | 0.025 | -0.007 | -0.014 | 0.027 | 0.026 |
| | distMult | 0.363 | 0.282 | 0.440 | 0.517 | 0.475 | 0.529 | 0.531 | 0.530 | 0.532 | 0.272 | 0.436 | 0.531 | 0.537 |
| PFAM_HS1 | ResnikBMA | 0.585 | 0.496 | 0.553 | 0.653 | 0.592 | 0.675 | 0.655 | 0.709 | 0.730 | **0.588** | 0.677 | 0.655 | 0.686 |
| | SimGIC | 0.546 | 0.557 | 0.600 | 0.649 | 0.616 | **0.679** | 0.655 | **0.712** | **0.733** | 0.576 | **0.684** | 0.655 | **0.691** |
| | RDF2Vec | 0.595 | 0.567 | 0.613 | 0.665 | 0.636 | 0.667 | **0.668** | 0.686 | 0.686 | 0.466 | 0.635 | **0.668** | 0.678 |
| | TransE | -0.096 | -0.091 | -0.059 | -0.129 | -0.104 | 0.120 | 0.131 | 0.124 | 0.126 | 0.024 | 0.042 | 0.131 | 0.131 |
| | distMult | 0.499 | 0.293 | 0.411 | 0.568 | 0.515 | 0.576 | 0.575 | 0.584 | 0.583 | 0.334 | 0.507 | 0.575 | 0.587 |

Table S5 Continued from previous page

| | | Static Similarity | | | | | Supervised Similarity | | | | | | | |
|---|---|---|---|---|---|---|---|---|---|---|---|---|---|---|
| | | **BP** | **CC** | **MF** | **AVG** | **MAX** | **GP** | **LR** | **XGB** | **RF** | **DT** | **KNN** | **BR** | **MLP** |
| PFAM_HS3 | ResnikBMA | 0.601 | 0.526 | 0.562 | 0.670 | 0.607 | **0.689** | 0.669 | **0.724** | **0.745** | **0.602** | **0.699** | 0.669 | **0.700** |
| | SimGIC | 0.554 | 0.575 | 0.611 | 0.659 | 0.622 | 0.687 | 0.665 | 0.721 | 0.742 | 0.596 | 0.694 | 0.665 | 0.698 |
| | RDF2Vec | 0.608 | 0.591 | 0.628 | 0.679 | 0.649 | 0.685 | **0.683** | 0.700 | 0.700 | 0.492 | 0.650 | **0.683** | 0.690 |
| | TransE | -0.106 | -0.100 | -0.073 | -0.144 | -0.111 | | 0.145 | 0.143 | 0.145 | 0.030 | 0.045 | 0.145 | 0.147 |
| | distMult | 0.062 | 0.304 | 0.423 | 0.433 | 0.371 | 0.478 | 0.478 | 0.483 | 0.477 | 0.229 | 0.379 | 0.478 | 0.488 |
| PFAM_SC1 | ResnikBMA | 0.466 | 0.365 | 0.535 | 0.569 | 0.556 | 0.609 | 0.588 | **0.763** | 0.687 | **0.640** | 0.677 | 0.588 | 0.666 |
| | SimGIC | 0.467 | 0.375 | 0.596 | 0.598 | 0.613 | **0.674** | 0.625 | 0.707 | **0.708** | 0.620 | **0.688** | 0.625 | **0.688** |
| | RDF2Vec | 0.464 | 0.366 | 0.611 | 0.582 | 0.614 | 0.652 | **0.630** | 0.681 | 0.675 | 0.479 | 0.634 | **0.630** | 0.663 |
| | TransE | 0.059 | 0.054 | 0.063 | 0.097 | 0.077 | 0.080 | 0.094 | 0.078 | 0.089 | -0.003 | 0.019 | 0.094 | 0.094 |
| | distMult | 0.422 | 0.272 | 0.419 | 0.522 | 0.490 | 0.530 | 0.527 | 0.541 | 0.544 | 0.284 | 0.450 | 0.527 | 0.545 |
| PFAM_SC3 | ResnikBMA | 0.508 | 0.394 | 0.557 | 0.596 | 0.587 | 0.645 | 0.603 | **0.692** | 0.668 | **0.639** | 0.684 | 0.603 | 0.676 |
| | SimGIC | 0.442 | 0.353 | 0.598 | 0.586 | 0.612 | **0.669** | 0.619 | 0.688 | **0.709** | 0.619 | **0.686** | 0.619 | **0.684** |
| | RDF2Vec | 0.478 | 0.394 | 0.607 | 0.592 | 0.615 | 0.665 | **0.631** | 0.671 | 0.680 | 0.464 | 0.626 | **0.631** | 0.674 |
| | TransE | 0.073 | 0.065 | 0.074 | 0.117 | 0.094 | 0.099 | 0.111 | 0.081 | 0.098 | 0.007 | 0.020 | 0.111 | 0.114 |
| | distMult | 0.450 | 0.275 | 0.433 | 0.545 | 0.504 | 0.563 | 0.552 | 0.554 | 0.567 | 0.311 | 0.479 | 0.552 | 0.572 |
| PFAM_EC1 | ResnikBMA | 0.364 | 0.339 | 0.373 | 0.454 | 0.425 | **0.530** | **0.446** | **0.578** | **0.600** | **0.506** | **0.564** | **0.446** | **0.542** |
| | SimGIC | 0.244 | 0.338 | 0.261 | 0.384 | 0.409 | 0.500 | 0.393 | 0.552 | 0.590 | 0.484 | 0.557 | 0.393 | 0.535 |
| | RDF2Vec | 0.244 | 0.338 | 0.260 | 0.383 | 0.377 | 0.422 | 0.388 | 0.434 | 0.442 | 0.219 | 0.382 | 0.388 | 0.424 |
| | TransE | -0.023 | 0.014 | -0.006 | -0.009 | -0.006 | | 0.002 | -0.006 | -0.013 | 0.014 | 0.025 | 0.003 | -0.015 |
| | distMult | 0.042 | 0.268 | 0.169 | 0.244 | 0.245 | 0.320 | 0.304 | 0.298 | 0.315 | 0.128 | 0.198 | 0.304 | 0.323 |
| PFAM_EC3 | ResnikBMA | 0.377 | 0.376 | 0.356 | 0.485 | 0.463 | **0.528** | **0.489** | 0.496 | 0.506 | **0.466** | **0.535** | **0.489** | **0.532** |
| | SimGIC | 0.296 | 0.366 | 0.229 | 0.418 | 0.433 | 0.494 | 0.433 | **0.552** | **0.507** | 0.440 | 0.499 | 0.433 | 0.495 |
| | RDF2Vec | 0.245 | 0.400 | 0.261 | 0.431 | 0.443 | 0.467 | 0.467 | 0.456 | 0.470 | 0.268 | 0.413 | 0.467 | 0.475 |
| | TransE | -0.007 | 0.024 | -0.050 | -0.019 | -0.026 | | 0.028 | 0.011 | -0.014 | 0.033 | -0.010 | 0.029 | 0.017 |
| | distMult | 0.099 | 0.251 | 0.127 | 0.249 | 0.225 | 0.273 | 0.269 | 0.246 | 0.268 | 0.125 | 0.179 | 0.269 | 0.272 |

Table S6: Pearson correlation coefficient between PPI and static similarity for the baselines and the median Pearson correlation coefficient between PPI and supervised similarity. Within each dataset, for each ML algorithm, the best SSM is in bold. The SSM is underlined when, using the same ML algorithm, there are no significant differences between the SSM and the best SSM (using $\alpha = 0.01$). Within each dataset, for each SSM, gray shading indicates the best ML algorithm. The light gray indicates that, using the same SSM, there are no significant differences between the ML algorithm and the best ML algorithm (using $\alpha = 0.01$).

| | | **Static Similarity** | | | | | **Supervised Similarity** | | | | | | | |
|---|---|---|---|---|---|---|---|---|---|---|---|---|---|---|
| | | **BP** | **CC** | **MF** | **AVG** | **MAX** | **GP** | **LR** | **XGB** | **RF** | **DT** | **KNN** | **BR** | **MLP** |
| PPI_ALL1 | ResnikBMA | 0.545 | 0.486 | 0.353 | 0.569 | 0.545 | **0.628** | **0.580** | **0.634** | **0.634** | **0.421** | **0.566** | **0.580** | **0.635** |
| | SimGIC | 0.486 | 0.428 | 0.318 | 0.496 | 0.458 | 0.578 | 0.514 | 0.584 | 0.585 | 0.354 | 0.504 | 0.514 | 0.584 |
| | RDF2Vec | 0.457 | 0.404 | 0.353 | 0.477 | 0.440 | 0.500 | 0.488 | 0.510 | 0.505 | 0.256 | 0.414 | 0.488 | 0.511 |
| | TransE | 0.031 | 0.087 | 0.035 | 0.071 | 0.076 | 0.086 | 0.087 | 0.083 | 0.085 | 0.012 | 0.023 | 0.087 | 0.086 |
| | distMult | 0.452 | 0.244 | 0.015 | 0.388 | 0.396 | 0.493 | 0.471 | 0.504 | 0.504 | 0.247 | 0.403 | 0.471 | 0.505 |
| PPI_ALL3 | ResnikBMA | 0.523 | 0.446 | 0.330 | 0.545 | 0.507 | **0.603** | **0.555** | **0.607** | **0.607** | **0.382** | **0.531** | **0.555** | **0.607** |
| | SimGIC | 0.457 | 0.390 | 0.290 | 0.462 | 0.416 | 0.543 | 0.481 | 0.548 | 0.541 | 0.307 | 0.467 | 0.481 | 0.549 |
| | RDF2Vec | 0.420 | 0.352 | 0.314 | 0.433 | 0.395 | 0.454 | 0.444 | 0.462 | 0.458 | 0.205 | 0.357 | 0.444 | 0.462 |
| | TransE | 0.043 | 0.098 | 0.040 | 0.082 | 0.089 | 0.089 | 0.099 | 0.090 | 0.095 | 0.008 | 0.027 | 0.099 | 0.098 |
| | distMult | 0.005 | 0.213 | 0.019 | 0.127 | 0.129 | 0.213 | 0.209 | 0.216 | 0.214 | 0.044 | 0.110 | 0.209 | 0.221 |
| PPI_DM1 | ResnikBMA | 0.789 | 0.765 | 0.597 | 0.812 | 0.796 | **0.890** | **0.829** | **0.869** | **0.875** | **0.768** | **0.913** | **0.829** | **0.847** |
| | SimGIC | 0.682 | 0.689 | 0.530 | 0.705 | 0.727 | 0.861 | 0.750 | 0.855 | 0.845 | 0.710 | 0.828 | 0.750 | 0.825 |
| | RDF2Vec | 0.659 | 0.721 | 0.560 | 0.724 | 0.734 | 0.773 | 0.751 | 0.782 | 0.787 | 0.654 | 0.746 | 0.751 | 0.748 |
| | TransE | 0.069 | 0.082 | 0.134 | 0.157 | 0.135 | 0.091 | 0.135 | 0.089 | 0.163 | 0.045 | 0.127 | 0.139 | 0.143 |
| | distMult | 0.739 | 0.551 | 0.492 | 0.781 | 0.739 | 0.835 | 0.802 | 0.815 | 0.792 | 0.703 | 0.787 | 0.802 | 0.816 |
| PPI_DM3 | ResnikBMA | 0.765 | 0.720 | 0.536 | 0.782 | 0.762 | **0.854** | 0.785 | **0.819** | **0.869** | **0.772** | **0.902** | 0.784 | 0.795 |
| | SimGIC | 0.650 | 0.643 | 0.483 | 0.652 | 0.675 | 0.828 | 0.677 | 0.818 | 0.818 | 0.730 | 0.809 | 0.679 | **0.807** |
| | RDF2Vec | 0.607 | 0.673 | 0.524 | 0.673 | 0.683 | 0.679 | 0.674 | 0.681 | 0.669 | 0.505 | 0.677 | 0.675 | 0.681 |
| | TransE | 0.094 | 0.129 | 0.131 | 0.196 | 0.199 | 0.199 | 0.181 | 0.170 | 0.198 | -0.004 | 0.140 | 0.183 | 0.211 |
| | distMult | 0.743 | 0.510 | 0.464 | 0.769 | 0.728 | 0.775 | **0.793** | 0.774 | 0.730 | 0.647 | 0.777 | **0.793** | 0.806 |
| PPI_HS1 | ResnikBMA | 0.465 | 0.404 | 0.347 | 0.501 | 0.454 | **0.549** | **0.509** | **0.560** | **0.560** | **0.311** | **0.473** | **0.509** | **0.561** |
| | SimGIC | 0.400 | 0.361 | 0.286 | 0.419 | 0.383 | 0.488 | 0.430 | 0.498 | 0.498 | 0.254 | 0.404 | 0.430 | 0.499 |
| | RDF2Vec | 0.369 | 0.295 | 0.280 | 0.390 | 0.359 | 0.403 | 0.401 | 0.418 | 0.412 | 0.175 | 0.306 | 0.401 | 0.419 |
| | TransE | 0.053 | 0.066 | 0.114 | 0.101 | 0.107 | 0.106 | 0.117 | 0.108 | 0.116 | 0.032 | 0.048 | 0.117 | 0.118 |
| | distMult | 0.009 | 0.184 | 0.177 | 0.191 | 0.174 | 0.240 | 0.239 | 0.239 | 0.241 | 0.065 | 0.117 | 0.239 | 0.245 |

Table S6 Continued from previous page

| | | Static Similarity | | | | | Supervised Similarity | | | | | | | |
|---|---|---|---|---|---|---|---|---|---|---|---|---|---|---|
| | | **BP** | **CC** | **MF** | **AVG** | **MAX** | **GP** | **LR** | **XGB** | **RF** | **DT** | **KNN** | **BR** | **MLP** |
| PPI_HS3 | ResnikBMA | 0.467 | 0.403 | 0.349 | 0.502 | 0.454 | **0.554** | **0.510** | **0.560** | **0.559** | **0.318** | **0.469** | **0.510** | **0.560** |
| | SimGIC | 0.399 | 0.358 | 0.285 | 0.417 | 0.381 | 0.495 | 0.430 | 0.505 | 0.504 | 0.254 | 0.406 | 0.430 | 0.503 |
| | RDF2Vec | 0.370 | 0.293 | 0.282 | 0.387 | 0.358 | 0.404 | 0.397 | 0.412 | 0.408 | 0.167 | 0.305 | 0.397 | 0.413 |
| | TransE | 0.054 | 0.068 | 0.113 | 0.100 | 0.107 | 0.105 | 0.115 | 0.110 | 0.115 | 0.020 | 0.042 | 0.115 | 0.114 |
| | distMult | 0.011 | 0.182 | 0.175 | 0.189 | 0.173 | 0.229 | 0.233 | 0.231 | 0.236 | 0.057 | 0.115 | 0.233 | 0.239 |
| PPI_SC1 | ResnikBMA | 0.621 | 0.563 | 0.401 | 0.643 | 0.639 | **0.701** | **0.658** | **0.714** | **0.712** | **0.540** | **0.663** | **0.658** | **0.712** |
| | SimGIC | 0.571 | 0.492 | 0.341 | 0.570 | 0.533 | 0.661 | 0.598 | 0.681 | 0.678 | 0.477 | 0.617 | 0.598 | 0.672 |
| | RDF2Vec | 0.563 | 0.512 | 0.419 | 0.594 | 0.545 | 0.634 | 0.603 | 0.644 | 0.644 | 0.417 | 0.569 | 0.603 | 0.644 |
| | TransE | 0.100 | 0.088 | 0.058 | 0.131 | 0.108 | 0.126 | 0.134 | 0.126 | 0.126 | 0.015 | 0.043 | 0.134 | 0.135 |
| | distMult | 0.436 | 0.387 | 0.288 | 0.531 | 0.483 | 0.560 | 0.538 | 0.570 | 0.567 | 0.325 | 0.475 | 0.538 | 0.571 |
| PPI_SC3 | ResnikBMA | 0.610 | 0.512 | 0.349 | 0.623 | 0.601 | **0.683** | **0.641** | **0.689** | **0.689** | **0.498** | **0.630** | **0.641** | **0.689** |
| | SimGIC | 0.548 | 0.440 | 0.294 | 0.532 | 0.479 | 0.634 | 0.572 | 0.644 | 0.645 | 0.426 | 0.580 | 0.572 | 0.636 |
| | RDF2Vec | 0.523 | 0.448 | 0.359 | 0.543 | 0.484 | 0.584 | 0.558 | 0.590 | 0.593 | 0.352 | 0.514 | 0.558 | 0.591 |
| | TransE | 0.128 | 0.097 | 0.058 | 0.151 | 0.124 | 0.150 | 0.167 | 0.126 | 0.156 | 0.032 | 0.071 | 0.167 | 0.167 |
| | distMult | 0.423 | 0.357 | 0.237 | 0.495 | 0.458 | 0.521 | 0.505 | 0.526 | 0.532 | 0.279 | 0.434 | 0.505 | 0.537 |
| PPI_EC1 | ResnikBMA | 0.613 | 0.647 | 0.496 | 0.703 | 0.707 | **0.730** | **0.707** | **0.751** | **0.741** | **0.626** | **0.711** | **0.707** | **0.721** |
| | SimGIC | 0.542 | 0.626 | 0.434 | 0.665 | 0.680 | 0.697 | 0.653 | 0.729 | 0.719 | 0.601 | 0.703 | 0.653 | 0.680 |
| | RDF2Vec | 0.558 | 0.584 | 0.501 | 0.682 | 0.641 | 0.702 | 0.672 | 0.684 | 0.691 | 0.532 | 0.645 | 0.672 | 0.672 |
| | TransE | 0.086 | 0.043 | -0.030 | 0.055 | 0.022 | 0.029 | 0.061 | 0.015 | 0.030 | 0.012 | -0.027 | 0.061 | 0.046 |
| | distMult | 0.336 | 0.463 | 0.295 | 0.531 | 0.470 | 0.545 | 0.545 | 0.581 | 0.543 | 0.394 | 0.472 | 0.545 | 0.555 |
| PPI_EC3 | ResnikBMA | 0.568 | 0.621 | 0.438 | 0.678 | 0.673 | **0.683** | **0.692** | **0.716** | **0.705** | **0.586** | 0.687 | **0.692** | **0.701** |
| | SimGIC | 0.516 | 0.604 | 0.388 | 0.641 | 0.649 | 0.652 | 0.658 | 0.631 | 0.652 | 0.531 | **0.704** | 0.658 | 0.664 |
| | RDF2Vec | 0.562 | 0.539 | 0.458 | 0.656 | 0.599 | 0.650 | 0.647 | 0.641 | 0.662 | 0.472 | 0.645 | 0.647 | 0.643 |
| | TransE | 0.155 | 0.104 | 0.002 | 0.144 | 0.087 | 0.093 | 0.124 | 0.101 | 0.096 | -0.013 | -0.063 | 0.129 | 0.134 |
| | distMult | 0.340 | 0.457 | 0.264 | 0.522 | 0.456 | 0.525 | 0.515 | 0.509 | 0.498 | 0.333 | 0.511 | 0.515 | 0.506 |

Table S7: Pearson correlation coefficient between $\mathrm{Sim}_{\mathrm{PS}}$ and static similarity for the baselines and the median Pearson correlation coefficient between $\mathrm{Sim}_{\mathrm{PS}}$ and supervised similarity for gene dataset. Within each dataset, for each ML algorithm, the best SSM is in bold. The SSM is underlined when, using the same ML algorithm, there are no significant differences between the SSM and the best SSM (using $\alpha = 0.01$). Within each dataset, for each SSM, gray shading indicates the best ML algorithm. The light gray indicates that, using the same SSM, there are no significant differences between the ML algorithm and the best ML algorithm (using $\alpha = 0.01$).

| | **Static Similarity** | | | | | | **Supervised Similarity** | | | | | | | |
|---|---|---|---|---|---|---|---|---|---|---|---|---|---|---|
| | **HPO** | **BP** | **CC** | **MF** | **AVG** | **MAX** | **GP** | **LR** | **XGB** | **RF** | **DT** | **KNN** | **BR** | **MLP** |
| ResnikBMA | 0.601 | 0.210 | 0.142 | 0.055 | 0.413 | 0.552 | 0.611 | **0.605** | **0.648** | **0.648** | **0.414** | **0.569** | **0.605** | **0.636** |
| SimGIC | 0.489 | 0.205 | 0.158 | 0.095 | 0.399 | 0.429 | **0.625** | 0.495 | 0.630 | 0.629 | 0.397 | 0.546 | 0.495 | 0.632 |
| RDF2Vec | 0.526 | 0.230 | 0.182 | 0.123 | 0.396 | 0.351 | 0.550 | 0.531 | 0.563 | 0.564 | 0.334 | 0.487 | 0.531 | 0.559 |
| TransE | 0.042 | -0.019 | -0.012 | -0.019 | -0.004 | 0.016 | | 0.041 | 0.047 | 0.056 | 0.007 | 0.004 | 0.041 | 0.044 |
| distMult | 0.015 | 0.184 | 0.105 | 0.041 | 0.179 | 0.182 | 0.196 | 0.206 | 0.172 | 0.212 | 0.039 | 0.051 | 0.206 | 0.215 |
